# Chromatin-encoded IRX4 identity erosion bounds cardiomyocyte renewal in the infarcted human heart

**DOI:** 10.64898/2026.10.20.760578

**Authors:** Shigang Qiao, Lingmei Li, Chunyan He, Wei Li

## Abstract

Myocardial infarction leaves the adult human heart unable to regenerate, yet whether injured cardiomyocytes revert to an immature, proliferative state is unresolved. We integrated 1.59 million single-cell and single-nucleus profiles with single-nucleus chromatin accessibility, spatial transcriptomics, plasma-proteome Mendelian randomization and an independent cardiomyopathy cohort of 79 donors to define the injured cardiomyocyte state. Infarcted cardiomyocytes showed a module-wide erosion of ventricular identity, including IRX4, with stress-gene induction and a chromatin-encoded loss of the identity module (−0.69 standard units; P = 1.2 × 10□□). Proliferation was not restored: nuclei co-detecting two cell-cycle transcripts remained at 0.011% (5/46,200), and the mitotic signal was confined to the ischaemic core. Genetic dissection separated the identity node from its secretory output: no variant at the IRX4 locus reached genome-wide significance across eight phenotypes, whereas genetically predicted NT-proBNP was causally associated with heart failure (odds ratio 1.12 per standard deviation; 95% confidence interval 1.05–1.19). The ligand and its endothelial receptor NPR1 were spatially uncoupled, and IRX4 erosion did not generalize to chronic cardiomyopathy. The injured cardiomyocyte therefore adopts a chromatin-encoded, non-regenerative stress identity rather than a fetal one, and the natriuretic peptide that reports this state lies on a causal path to heart failure.

## Introduction

Heart failure after myocardial infarction remains a leading cause of death because adult cardiomyocytes cannot replace the myocardium lost to ischaemia[1–3]. Resident progenitors contribute negligibly to new cardiomyocytes[4], and the renewal rate of the adult human heart, approximately 1% per year, is insufficient for repair[2,3]. Regenerative strategies have therefore focused on re-engaging the cell-cycle machinery that is withdrawn during postnatal maturation, when a glycolytic programme gives way to fatty-acid oxidation[5–7].

The conceptual anchor of this field is dedifferentiation, the idea that injury returns cardiomyocytes to an immature, proliferative state. Forced expression of ERBB2, cell-cycle regulators, Hippo pathway components or microRNAs can re-initiate proliferation and, in some models, produce functional repair[8–14]. Whether pathophysiological injury engages the same programme is a separate question. Injured cardiomyocytes re-express NPPA, NPPB and MYH6, the genes that define the fetal ventricle, and this fetal gene programme has been read as evidence of reversion[15]. The inference is not secure: the same genes are atrial-enriched stress markers, and their induction does not require loss of ventricular identity or re-entry into the cell cycle.

Four gaps have kept the question open. First, identity erosion has been inferred from labile, depth-sensitive transcript abundance, whereas chromatin, where the state should be fixed, has not been interrogated. Second, the genetic architecture of candidate identity factors has not been separated from that of their secreted markers, so it is unclear whether an identity node is itself a disease gene. Third, communication between the injured cardiomyocyte and its niche has been analysed as a network property without spatial constraints, conflating potential with realized signalling[16–19]. Fourth, most single-cell descriptions of myocardial infarction derive from one cohort, leaving unclear whether the state is specific to ischaemic injury or a general property of failing myocardium[20–23].

We addressed these gaps by combining a cross-species maturation reference, chromatin accessibility, spatial transcriptomics, a two-sample Mendelian randomization framework built on plasma protein quantitative trait loci, and an independent cardiomyopathy cohort. The results define a state that is chromatin-encoded, proliferatively silent, spatially specified to the injured zone, and genetically dissociable from the natriuretic biomarker that reports it.

## Results

### A multi-scale atlas resolves the identity of the stressed cardiomyocyte

We assembled a multi-scale reference spanning six public datasets and approximately 1.59 million cells and nuclei from human and mouse development, the adult human heart and myocardial infarction, integrated with single-nucleus ATAC-seq and Visium spatial transcriptomics (Figure 1A,B). The Human Cell Atlas cardiac-niche collection (D7; 704,296 cells) resolved 76 cell states across eight anatomical regions (Figure 1C,D), including twelve ventricular and atrial cardiomyocyte substates (Figure 1E) distinguished by the ventricular identity factor IRX4 and the stress markers NPPA and NPPB (Figure 1F). The distinction matters because the coarse cardiomyocyte category conceals a stress substate that already carries the molecular signature of the injured heart. Dataset composition, cell counts and modality are summarised in Supplementary Table S1 and Supplementary Figure S1.

**Figure 1.**
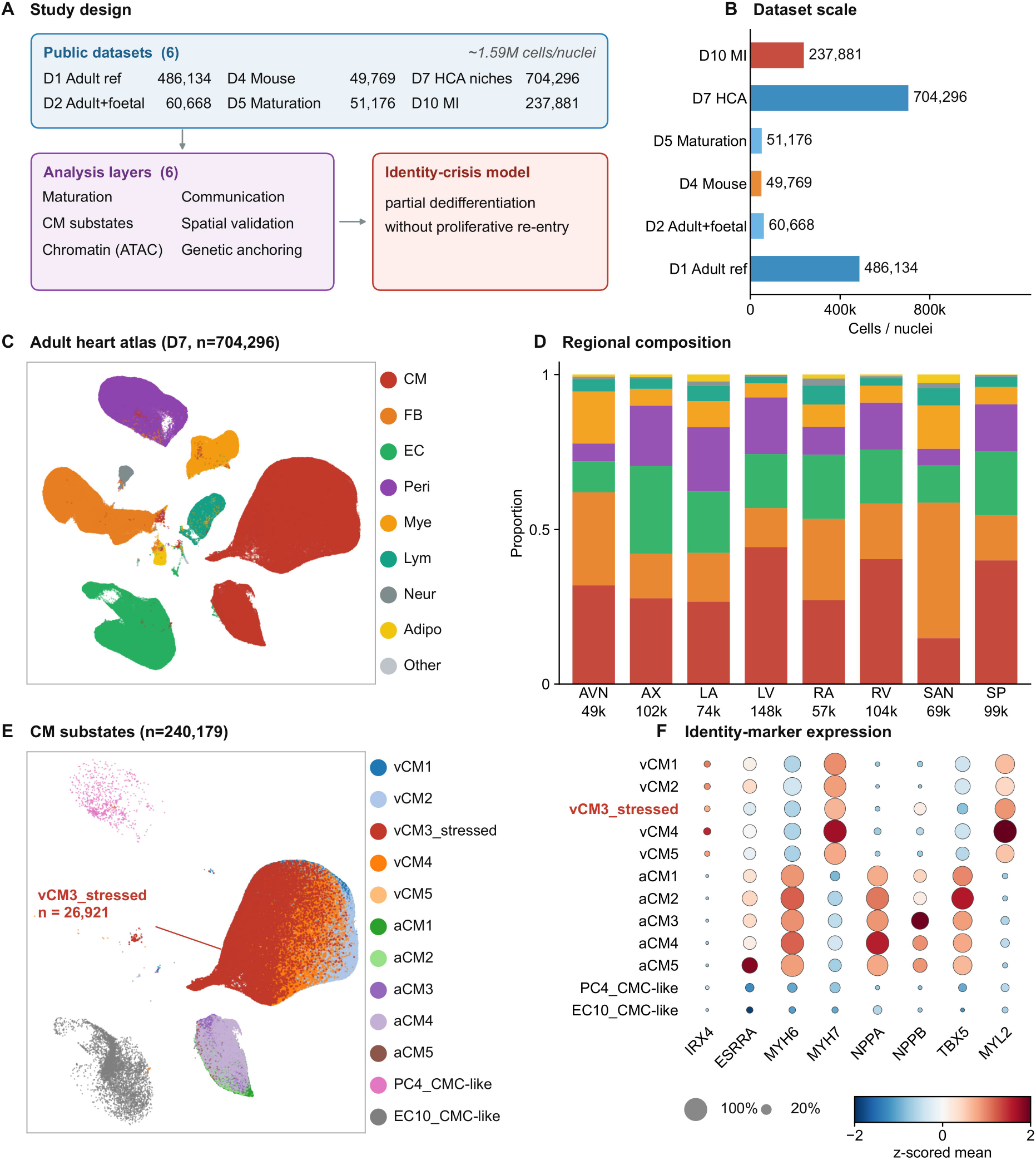
A multi-scale atlas of cardiac maturation, identity and communication. A, Study design. Six public datasets (approximately 1.59 million cells and nuclei) integrated across six analysis layers: maturation trajectory, cardiomyocyte substates, chromatin accessibility, intercellular communication, spatial validation and genetic context. B, Cells and nuclei per dataset and modality. C, Uniform manifold approximation and projection of the 704,296 cells of the adult human heart niche atlas (D7), coloured by cell class. D, Anatomical regions represented in D7. E, Twelve cardiomyocyte substates, including the stressed ventricular substate vCM3_stressed (n = 26,921). F, IRX4 and NPPB expression across cardiomyocyte substates; dot size, fraction of nuclei expressing; colour, mean log-normalised expression. Cell counts and composition are given in Supplementary Table S1 and Supplementary Figure S1. **Alt text:** Six-panel composite. A, schematic of the study design linking six public datasets to six analysis layers; B, cells or nuclei per dataset; C, UMAP of 704,296 adult heart cells coloured by cell class; D, anatomical regions represented; E, twelve cardiomyocyte substates; F, dot plot of IRX4 and NPPB expression across substates.

### A conserved cross-species maturation trajectory defines the non-regenerative baseline

The capacity of mammalian cardiomyocytes to divide is lost during the first postnatal weeks, in step with the metabolic switch from glycolysis to fatty-acid oxidation[5–7]. We reconstructed this trajectory from 38,295 human and 17,047 mouse cardiomyocytes spanning fetal-to-adult development (Figure 2A,B). Five programmes were scored by developmental maturity. Proliferation declined monotonically with maturation in both species (human Spearman ρ = −0.79, P = 0.036; mouse ρ = −0.83, P = 0.042), as did glycolysis in mouse (ρ = −0.83, P = 0.042). By contrast, fatty-acid oxidation (mouse ρ = +0.77, P = 0.072) and sarcomeric maturation markers (human ρ = +0.79, P = 0.036) rose (Figure 2C). At the gene level the switch reflected reciprocal declines in MKI67 and TOP2A and in glycolytic enzymes, with rises in CPT1B, PDK4, MYL2 and TNNI3 (Figure 2D). Mechanistically, the oxygen-rich postnatal environment imposes DNA damage that enforces quiescence[24], and the oxidative state silences the proliferative network epigenetically[5,6]. The mature cardiomyocyte is therefore held in a configuration incompatible with division, not merely withdrawn from it.

**Figure 2.**
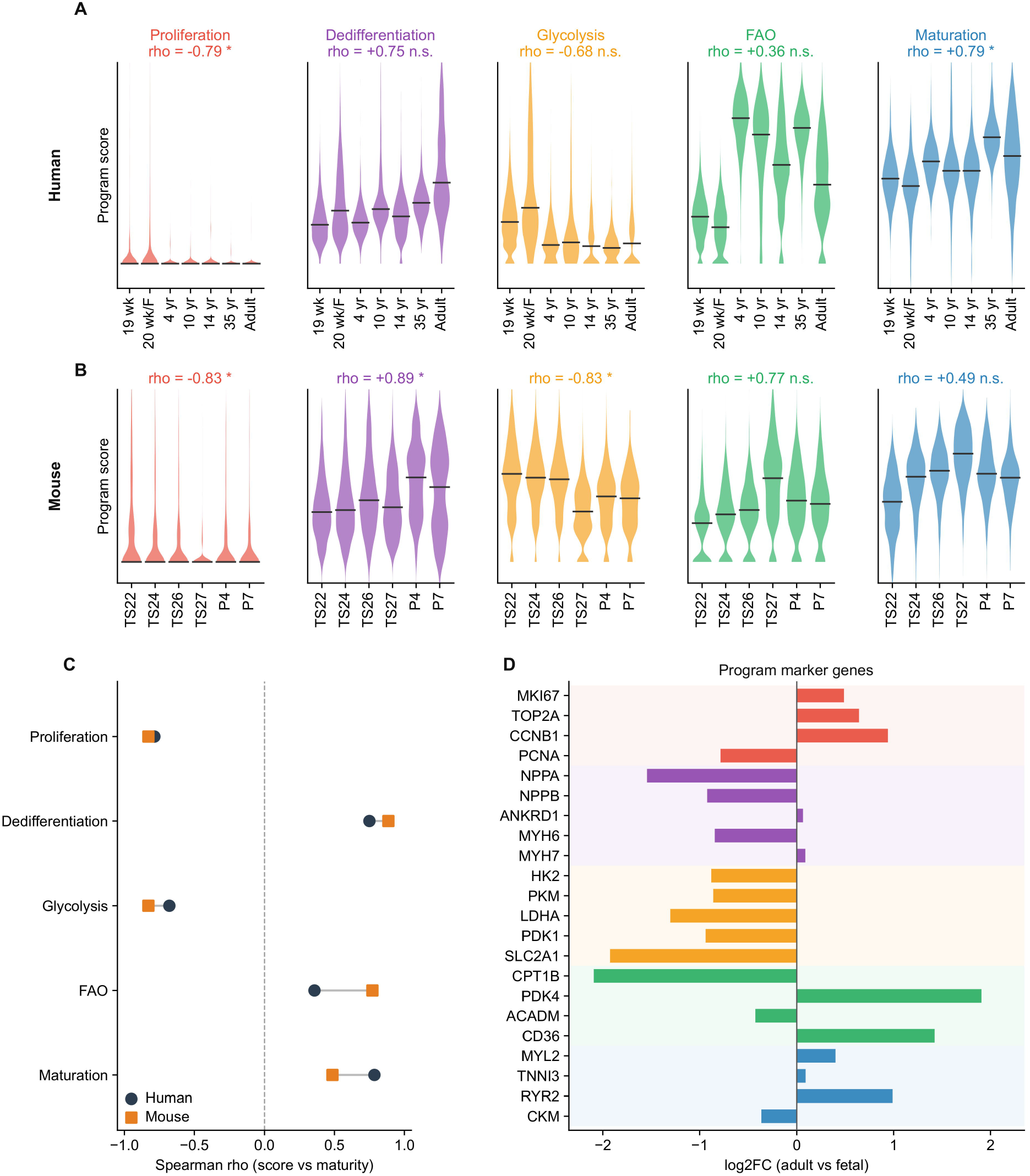
A conserved cross-species maturation trajectory defines the non-regenerative baseline. A, Programme scores (proliferation, dedifferentiation, glycolysis, fatty-acid oxidation and maturation) across seven human maturity stages (19 weeks of gestation to adult; n = 37,417 cardiomyocytes with an assigned stage); Spearman ρ versus maturity is given above each panel. B, The same programmes across six mouse stages (TS22 to P7; n = 17,047). C, Stage-mean programme scores with fitted trends; ρ and P values are from Spearman correlation across stage means. D, Gene-level expression of programme markers along the human maturity axis. The dedifferentiation programme does not decline with maturation, indicating that NPPA, NPPB and MYH6 behave as stress markers rather than as indices of immaturity. Complete scores are given in Supplementary Table S2 and gene-level heat maps in Supplementary Figure S2. **Alt text:** Four-panel figure. A and B, proliferation, dedifferentiation, glycolysis, fatty-acid oxidation and maturation scores across seven human and six mouse developmental stages; C, stage-mean scores with fitted trends and Spearman statistics; D, gene-level heat map of programme markers along human maturation.

The behaviour of the canonical dedifferentiation programme was informative. NPPA, NPPB and MYH6 did not decline with maturation; their composite score rose (human ρ = +0.75). These fetal genes therefore behave as atrial-enriched stress markers, not as faithful indices of immaturity, a distinction that has confounded the field and that motivated our use of transcription-factor activity to define the injured state. Gene-level programme expression is shown in Supplementary Figure S2A,B and complete scores in Supplementary Table S2.

### A stressed cardiomyocyte substate with eroded identity in the healthy heart

If injury-induced dedifferentiation were reversion, a stress substate present in healthy myocardium should carry fetal and proliferative markers. It did not. Sub-clustering resolved twelve cardiomyocyte substates, including a stressed ventricular population (vCM3_stressed, n = 26,921; Figure 3A). IRX4 was lower in vCM3_stressed than in resting ventricular cardiomyocytes (pseudobulk 9.5 versus 11.3 copies per million; ratio 0.84, 95% confidence interval 0.81–0.99; detectable transcript in 4.9% versus 5.9% of nuclei; 1,390 counts per 146.1 million versus 8,396 per 740.0 million; Figure 3B and Supplementary Table S3). ESRRA, a metabolic maturation factor, was also lower (Figure 3C), whereas NPPB was elevated 10.7-fold over pooled resting cardiomyocytes (Figure 3D). The identity–stress axis separated the stressed substate from its neighbours (Figure 3E).

**Figure 3.**
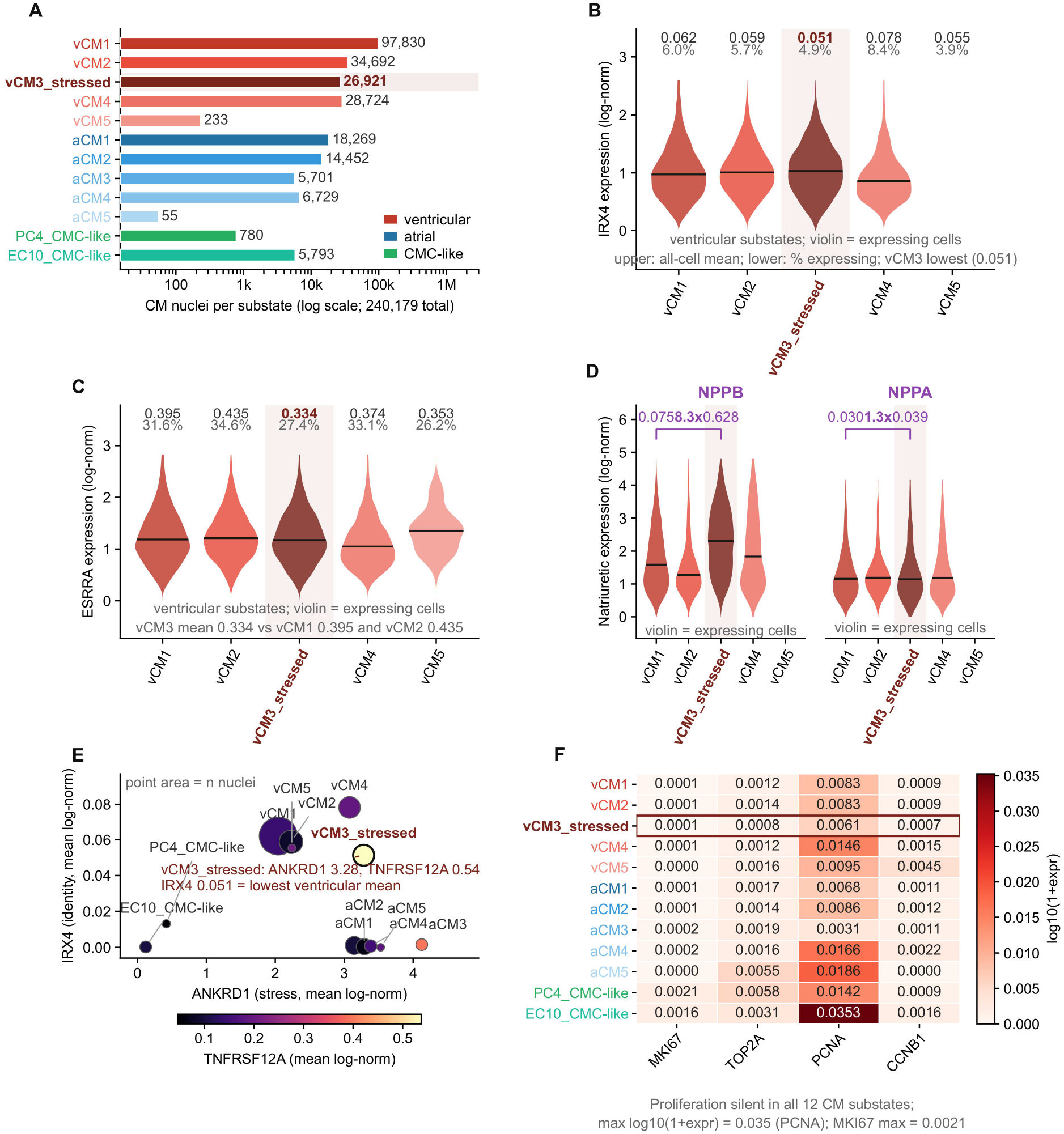
A stressed cardiomyocyte substate with eroded identity in the healthy heart. A, Composition of the twelve cardiomyocyte substates (D7; logarithmic scale). B, IRX4 across the five ventricular substates; values above the violins give mean log-normalised expression and the percentage of nuclei with a detectable transcript; vCM3_stressed is shaded. C, ESRRA across the same substates. D, NPPB and NPPA across substates. E, Identity–stress axis (IRX4 versus ANKRD1) separating the stressed substate from its resting neighbours. F, Proliferation score across all twelve substates, at the detection floor (maximum log□□(1 + expression) = 0.035). Horizontal bars mark the median; n = 26,921 nuclei for vCM3_stressed. Donor-level tests and marker correlations are given in Supplementary Table S3; substate composition in Supplementary Figure S3 and identity-gene expression in Supplementary Figure S4. **Alt text:** Six-panel figure. A, composition of the twelve cardiomyocyte substates; B, IRX4 expression across ventricular substates; C, ESRRA expression; D, NPPB and NPPA expression; E, identity-stress scatter (IRX4 versus ANKRD1); F, proliferation score across substates.

Two donor-level tests condition on different features. A depth-stratified comparison was significant (Δ = −0.010 ± 0.002, P = 0.002; leave-one-out −0.0100 [−0.0129, −0.0076]), whereas the unadjusted per-donor ratio was not (median 0.95, paired P = 0.067), because per-donor copies per million are imprecise when about 5% of nuclei are positive. In a donor-aware mixed-effects model with sequencing depth as a covariate, the reduction persisted (β = −0.007, 95% confidence interval −0.011 to −0.003; n = 159,443 ventricular cells), although it was more conservative under donor-clustered robust standard errors (P = 0.054 versus P = 4 × 10□□ with model-based standard errors). We therefore describe the transcript-level difference as modest and base inference on its agreement with transcription-factor activity and chromatin accessibility. Proliferation signals remained at the detection floor across all twelve substates (maximum log□□(1 + expression) = 0.035; Figure 3F).

Although vCM3_stressed expressed the atrial-enriched stress genes NPPA and NPPB, its marker profile across a 13-gene panel remained ventricular: it correlated with ventricular substates (r = 0.92–0.94) and not with atrial substates (r = 0.20–0.32; Supplementary Table S3), arguing against atrial conversion. We call this combination of lineage-identity erosion and stress-gene activation, without proliferative re-entry, an identity crisis. The state exists in healthy myocardium, with two implications. First, fetal gene re-expression after injury, the clinical biomarker of cardiac stress, does not denote a return to immaturity but entry into a proliferatively arrested stress substate. Second, infarction may expand a pre-existing, constitutively non-regenerative state rather than induce dedifferentiation, which explains the long-standing paradox of natriuretic-peptide elevation without myocardial regeneration. Substates are shown in Supplementary Figure S3 and identity-gene expression in Supplementary Figure S4.

### Infarction erodes identity without re-engaging proliferation

We projected infarct-zone cardiomyocytes (46,200 cells) onto this framework. If injury induced full dedifferentiation, these cells should shift toward the fetal quadrant and re-acquire proliferative capacity. Instead they showed partial identity erosion (Figure 4A). The myosin isoform ratio shifted toward MYH6 dominance (median 1.00 in infarction versus 0.43 in controls and 0.65 in adult reference hearts; donor-level medians 0.73 versus 0.49; Figure 4B), and the maturation score declined from 2.306 to 2.063. The proliferation score did not re-engage (0.0078 versus 0.0087; cell-level P = 0.015, Δ = 9 × 10□□), and fatty-acid oxidation was further elevated (1.104 to 1.385; donor-level effect sizes in Supplementary Table S4). Infarcted cardiomyocytes therefore occupied the mature metabolic–proliferative quadrant, not the fetal one (Figure 4C,D).

**Figure 4.**
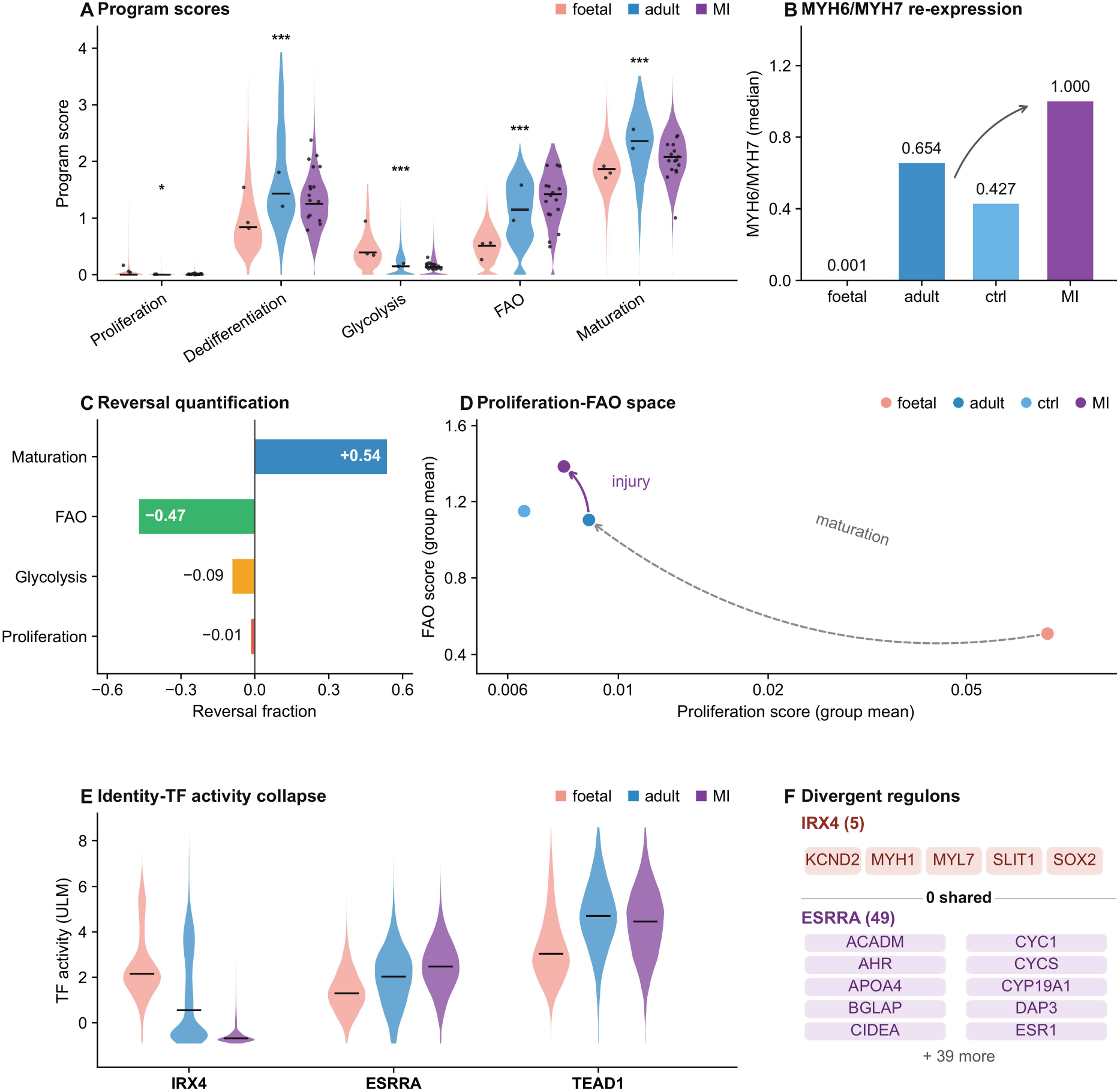
Infarction erodes identity without re-engaging proliferation. A, Programme scores of fetal (n = 22,994), adult (n = 7,547) and infarcted (n = 46,200) cardiomyocytes; violins show cells, black markers donor means, bars medians; asterisks denote donor-clustered tests (adult versus infarction; *P < 0.05). B, MYH6/MYH7 median ratio (fetal 0.001; adult 0.65; infarction 1.00; control 0.43). C, Reversal fraction of each programme relative to the fetal-to-adult span. D, Proliferation–fatty-acid oxidation biplot; infarcted cardiomyocytes occupy the mature metabolic quadrant. E, Double-positive (MKI67 and a second cell-cycle transcript) nuclei: 5 of 46,200 (0.011%) in infarction versus 0 of 18,310 in control. F, Transcription-factor activity for IRX4, ESRRA/ERRα and TEAD1 across fetal, adult and infarcted cardiomyocytes. Donor-level effect sizes are given in Supplementary Table S4, rare-event counts in Supplementary Figure S5, fetal transcription-factor switches in Supplementary Figure S6 and Supplementary Tables S5 and S6, and method checks in Supplementary Figure S7 and Supplementary Figure S8A–D. **Alt text:** Six-panel figure. A, programme scores in fetal, adult and infarcted cardiomyocytes; B, MYH6/MYH7 ratios; C, reversal fractions; D, proliferation-fatty-acid oxidation biplot; E, double-positive nuclei counts; F, IRX4, ESRRA/ERRalpha and TEAD1 activity.

Direct counting across 64,510 cardiomyocyte nuclei placed this state change against proliferative re-entry. Cohort-wide, MKI67-positive nuclei were detected at 0.24% in infarcted versus 0.31% in control myocardium (odds ratio 0.80, P = 0.17; donor-level P = 0.23). This comparison conceals a pronounced spatial structure: within the acutely ischaemic core, MKI67 detection rose to 1.13% [0.87–1.47], about 3.7-fold the control rate of 0.31% [0.24–0.40] (odds ratio 3.73; P = 2.7 × 10□¹¹; Supplementary Table S4). The border zone (0.14% [0.09–0.23]), remote myocardium (0.18% [0.13–0.24]) and fibrotic zone (0.03% [0.007–0.096]) showed no enrichment (Supplementary Figure S5A–D). This regional pattern agrees in direction with the increase in cardiomyocyte mitosis reported by immunostaining in human infarcted hearts[25], and it locates the response in tissue that is acutely deprived of oxygen, not across the injured ventricle.

The magnitude of the response is nonetheless bounded. Nuclei co-detecting two cell-cycle transcripts, the criterion that separates genuine cycling from single-transcript detection, remained at 0.011% in infarction (5 of 46,200; Wilson 95% confidence interval 0.005–0.025%) versus 0 of 18,310 in control (Figure 4E and Supplementary Figure S5D). No donor-level comparison of MKI67 detection was significant (P = 0.23), and the core signal was concentrated in a minority of donors and nuclei. Even if every double-positive nucleus completed division, the contribution would remain below 0.03% of the cardiomyocyte population. The frequencies are compatible with counts after human infarction[25] and with mouse dedifferentiation models[26], which is why we describe cardiomyocyte mitosis after infarction as regionally confined and quantitatively insufficient, not absent.

Transcription-factor activity resolved the regulatory logic. Using the CollecTRI regulon resource (1,185 factors) with univariate linear models across fetal, adult and infarcted cardiomyocytes, the fetal switches (KLF7, CREB3L1, IRF7, RXRA) were not reactivated (Supplementary Figure S6A,B). Instead the most prominent change was collapse of IRX4 activity (1.267 to −0.502; a five-target regulon, too small for independent regulon-level validation; activity units are dimensionless and not commensurable with transcript abundance), whereas the metabolic maturation factor ESRRA/ERRα rose (1.998 to 2.472; donor-level P = 0.10) and TEAD1 remained high (Figure 4F). This asymmetry is the key observation: the injured cardiomyocyte does not turn back the clock toward a fetal programme but loses a subset of mature-identity factors while strengthening its metabolic programme. Because the IRX4 regulon is too small for pathway-level inference, its centrality rests on convergence across transcript, chromatin and genetic layers. The complete activity matrix is in Supplementary Table S5, fetal switches in Supplementary Table S6, and method checks in Supplementary Figure S7 and Supplementary Figure S8A–D.

### Identity erosion is encoded in chromatin and is donor-robust

A transcriptional state is durable only if it is fixed in chromatin. We therefore examined single-nucleus ATAC-seq gene-activity scores across the infarcted human heart (46,086 nuclei; 21,923 cardiomyocyte-lineage nuclei from 20 donors; Figure 5A). IRX4 accessibility declined monotonically along the injury gradient, from 0.387 in control through 0.374 at the border and 0.279 in fibrotic tissue to 0.255 in the ischaemic core. In a donor-clustered model the ischaemic–control difference was −0.126 (95% confidence interval −0.190 to −0.062; P = 1.2 × 10□□), and the effect was stable in leave-one-donor-out analysis (median −0.132, range −0.151 to −0.111; n = 20 donors; Figure 5A,D). ESRRA (−0.177, P = 3.6 × 10□□) and MEF2C (−0.052, P = 0.077) moved in the same direction, whereas TEAD1 was largely retained (−0.081, P = 0.33; Figure 5B,C). The identity module as a whole fell by 0.694 standard units (P = 1.2 × 10□□), and chamber myosin loci lost accessibility (MYH7 −0.551, P = 9.5 × 10□□) while the stress-peptide locus NPPB gained it (+0.117, P = 0.29; Figure 5E). The response is therefore an injury-selective reconfiguration of the identity programme, not a global loss of accessibility.

**Figure 5.**
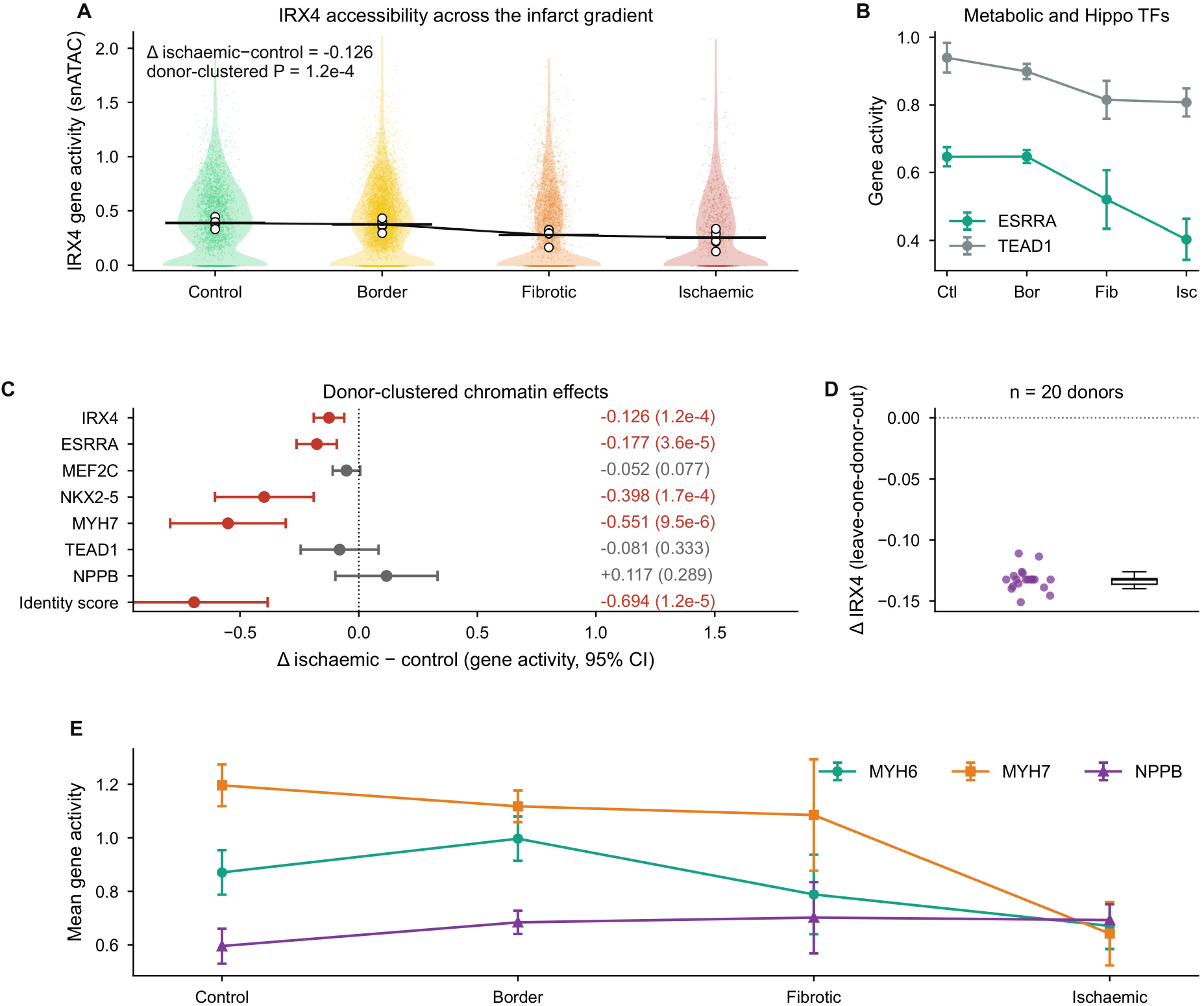
Identity erosion is encoded in chromatin and is donor-robust. A, IRX4 chromatin accessibility (single-nucleus ATAC gene activity) along the infarct gradient in 21,923 cardiomyocyte-lineage nuclei from 20 donors. Violins show nuclei, open circles donor means, the black line the donor-mean trajectory; the donor-clustered ischaemic–control difference is −0.126 (P = 1.2 × 10□□). B, ESRRA and TEAD1 accessibility across zones (mean ± s.e.m. of donor means). C, Donor-clustered ischaemic–control effects with 95% confidence intervals for eight readouts; red, P < 0.05. D, Leave-one-donor-out distribution of the IRX4 ischaemic–control difference (20 iterations; median −0.132, range −0.151 to −0.111). E, MYH6, MYH7 and NPPB accessibility across zones. Values are gene-activity scores; complete per-nucleus and per-donor values are given in Supplementary Tables S7 and S8, regulon enrichment in Supplementary Figure S10A,B, donor-level values in Supplementary Figure S12, and depth-confounding analyses in Supplementary Figure S9. **Alt text:** Five-panel figure. A, IRX4 chromatin accessibility along the infarct gradient; B, ESRRA and TEAD1 accessibility by zone; C, donor-clustered effect sizes for eight readouts; D, leave-one-donor-out distribution; E, MYH6, MYH7 and NPPB accessibility.

Two qualifications apply. First, the transcript-level decline of IRX4 is confounded by sequencing depth: ischaemic-zone nuclei had a fourfold lower median depth (2,570 versus 10,254 unique molecular identifiers), and every transcript examined fell in parallel (NKX2-5 33.3% to 7.0%; MYL2 89.0% to 34.9%; RPLP0 17.8% to 5.7%; Supplementary Figure S9A–C). After depth adjustment the reduction persisted (odds ratio 0.59, 95% confidence interval 0.47–0.74) but was no larger than that of other identity transcripts, so transcript detection alone does not establish selective IRX4 loss. Chromatin accessibility, which is depth-normalized by design, shows the same module-level pattern, not IRX4 selectivity; IRX4 is distinguished not by the magnitude of its change but by its position as the ventricular-specifying factor whose loss erodes the module in model systems[27–29]. Second, regulon enrichment across 750 CollecTRI factors (Supplementary Table S7) showed only negative trends for cardiac-identity regulators (TBX5 P = 0.019, ESRRA P = 0.039; TEAD1, NKX2-5 and MEF2C concordantly negative), none surviving Benjamini–Hochberg correction. The single factor passing correction was MYC (q = 4 × 10□□; 808 targets; mean target Δlog□ fold change −0.05), a modest contraction we interpret as a dampened proliferation–metabolism programme, not a causal node (Supplementary Figure S10A,B, Supplementary Figure S11A–D, Supplementary Table S7 and per-donor statistics in Supplementary Table S8). Per-donor zone means and paired contrasts are given in Supplementary Figure S12A–C. Chromatin accessibility therefore establishes that identity erosion is a durable, epigenetically encoded state that is unlikely to reverse spontaneously; a transcriptional epiphenomenon would not show this stability. The same data also expose a cross-layer divergence: ESRRA activity rose in infarction even as its locus lost accessibility, so activity and chromatin accessibility are not interchangeable readouts. The identity programme is most stringently defined by IRX4, whose erosion is concordant across transcript, chromatin and genetic layers, with the caveat that its regulon is too small to provide an independent pathway-level test.

### Genetic dissection separates the identity node from its natriuretic output

The analyses above are correlational. To ask whether the identity node or its secretory output is causally connected to disease, we turned to human genetics. We scanned the IRX4 locus (chr5:1.85–1.95 Mb, GRCh38) across eight FinnGen R13 phenotypes[30] in 608 common variants passing quality control (Supplementary Table S9 and Supplementary Figure S13A,B). No variant reached genome-wide significance in any phenotype (smallest P = 1.9 × 10□□ for dementia and 7.4 × 10 for heart failure; all phenotype minima between 1.9 × 10□□ and 6.3 × 10□³; Figure 6A,B). The lead conduction variant rs260399, which lies within the IRX4 gene body and is associated with QRS duration[31,32], showed no association with heart failure (odds ratio 0.99 per C allele, 95% confidence interval 0.98–1.01; P = 0.33) or with any other phenotype (all P > 0.15; Figure 6C). IRX4 is therefore not a common-variant disease gene for heart failure, despite being the chromatin-defined node of the injured cardiomyocyte.

**Figure 6.**
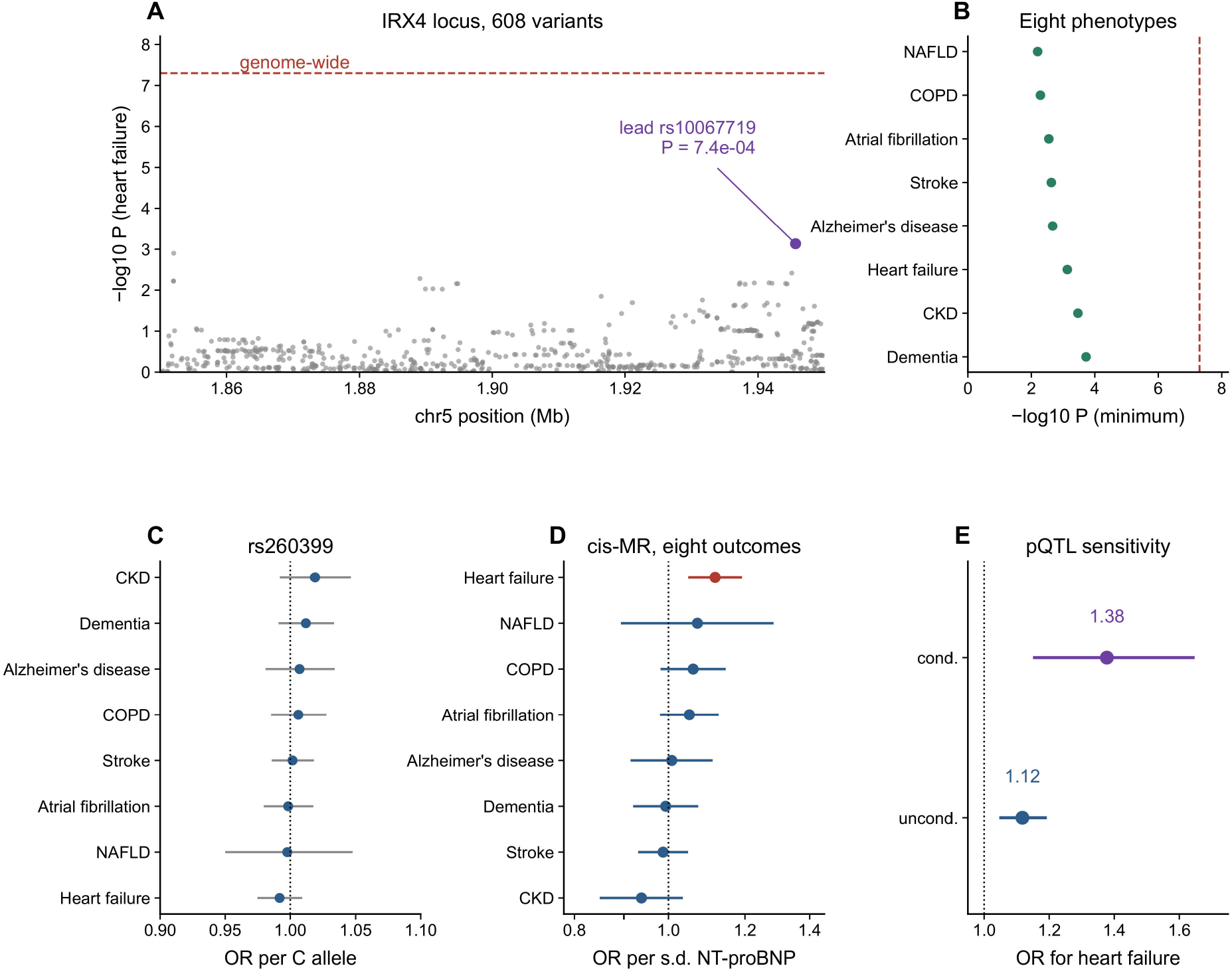
Genetic dissection separates the identity node from its natriuretic output. A, Regional association plot of the IRX4 locus (chr5:1.85–1.95 Mb, GRCh38) for heart failure in FinnGen release 13 (608 common variants); the dashed line marks genome-wide significance (5 × 10□□) and the purple point the lead variant. B, Phenome-wide scan of the same locus across eight phenotypes; no phenotype reaches genome-wide significance. C, Effect of the IRX4 lead conduction variant rs260399 on eight phenotypes (odds ratio per C allele with 95% confidence interval); all estimates are null. D, Two-sample Mendelian randomization of genetically predicted NT-proBNP (cis-pQTL rs198389; F = 984; R² = 3.2%) on the same eight phenotypes; heart failure is the only significant outcome (odds ratio 1.12 per s.d., 95% confidence interval 1.05–1.19; P = 3.7 × 10□□). E, Sensitivity of the estimates to the conditional (0.090) rather than the unconditional (0.259) pQTL effect. Full statistics for the genetic analyses are given in Supplementary Table S10. **Alt text:** Five-panel figure. A, regional association plot at the IRX4 locus for heart failure; B, phenome-wide scan across eight phenotypes; C, effects of rs260399; D, Mendelian randomization estimates for genetically predicted NT-proBNP; E, sensitivity to the conditional pQTL effect.

Its principal secreted product behaved differently. The cis-pQTL for NT-proBNP (rs198389; β = 0.259 s.d. per G allele; P = 5.7 × 10□²¹□; F = 984; R² = 3.2%)[33] provided a strong instrument for two-sample Mendelian randomization[34,35]. Genetically predicted NT-proBNP was associated with heart failure (odds ratio 1.12 per s.d., 95% confidence interval 1.05–1.19; P = 3.7 × 10□□; Supplementary Table S10). It was not associated with atrial fibrillation (1.05, 0.98–1.12; P = 0.15), stroke, dementia, Alzheimer’s disease, chronic kidney disease, chronic obstructive pulmonary disease or non-alcoholic fatty liver disease (all P > 0.12; Figure 6D and Supplementary Table S10). The estimate was robust to the use of the conditional rather than the unconditional pQTL effect (odds ratio 1.38, 1.15–1.64; Figure 6E). Three conclusions follow. First, the identity node and the biomarker are genetically separable: IRX4 erosion marks the cell state, but inherited variation at IRX4 does not confer heart-failure risk at common-variant frequencies. Second, the natriuretic output of that state is itself on a causal path to heart failure, which provides orthogonal, non-transcriptomic support for the secretory arm of the phenotype and does so without reliance on any single cohort. Third, the tissue specificity is consistent with the source: NPPB is heart-enriched (Open Targets genetic association with hypertensive disorder and blood pressure; Supplementary Figure S14A–E), whereas the other outcomes tested are not. The dissociation cautions against interpreting natriuretic-peptide elevation as a surrogate for cardiomyocyte identity: the two are linked within the cell but not across the germline.

### The intercellular layer does not resolve beyond permutation nulls

Cardiomyocyte identity is maintained, and its erosion sensed, within a cellular niche[36,37]. Using a ligand–receptor compendium (OmniPath; 4,349 expressed pairs[18]) across 447,885 cells of the adult human heart[20], we constructed a cell-type by cell-type communication network (Figure 7A). The network had a directional structure that undirected analyses conceal: fibroblasts were the most prominent senders (out-strength 176.1; net +59.5), whereas cardiomyocytes (net −17.3) and pericytes (net −25.0) were net receivers (Figure 7B,C). Endothelial cells were strongly bidirectional. Fibroblasts are the principal secretors of matrix, growth factors and cytokines that maintain the niche[36,38]; cardiomyocytes are largely recipients. This architecture provides the baseline against which injury can be measured.

**Figure 7.**
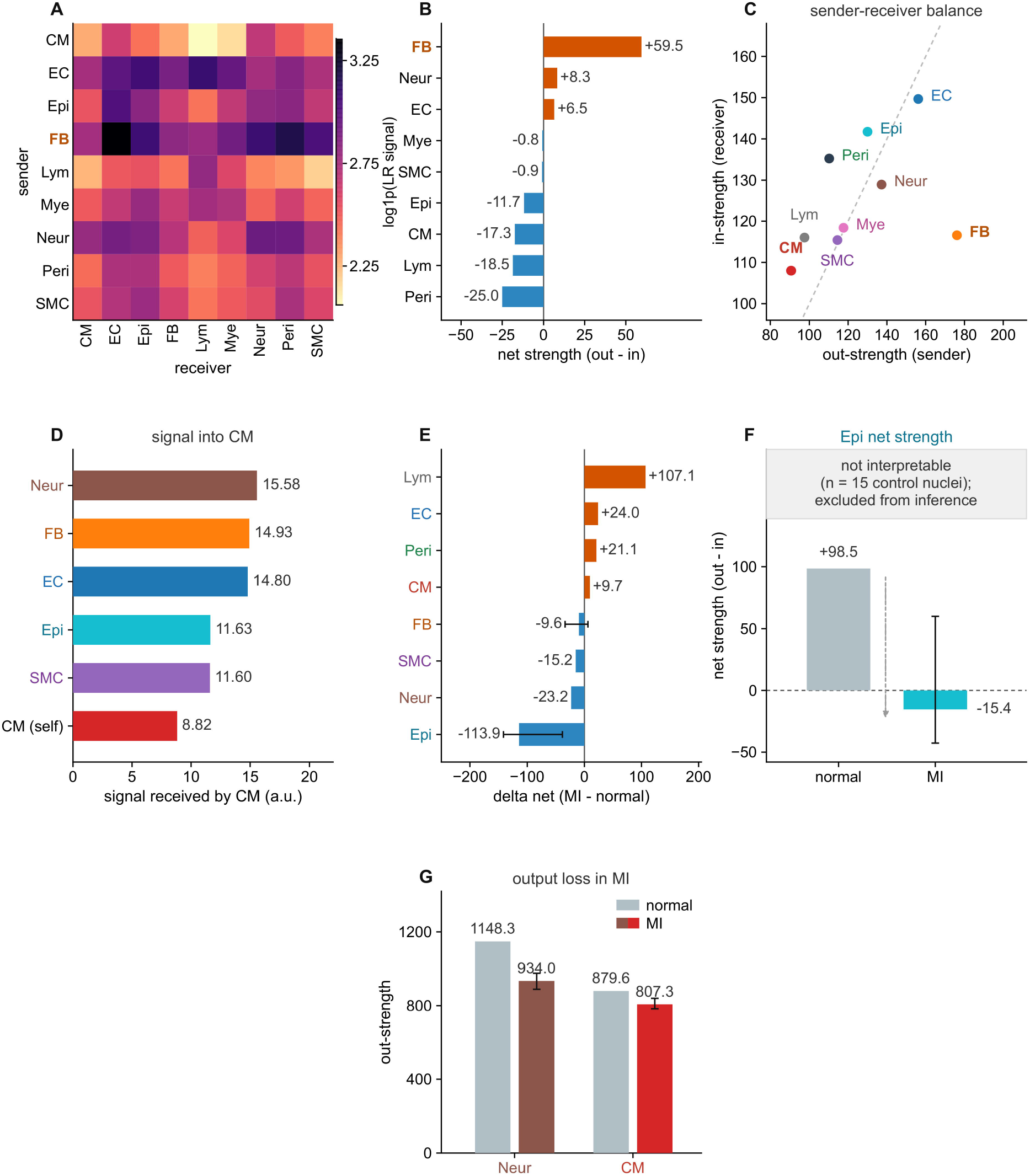
Directional architecture of cardiac intercellular communication and its limits after infarction. A, Cell-type by cell-type communication network (447,885 cells; OmniPath, 4,349 expressed pairs). B, Net strength (out minus in) for the nine classes in the D1 structural baseline; fibroblasts are the most prominent senders (net +59.5), cardiomyocytes (net −17.3) and pericytes (net −25.0) are net receivers. C, Ranked net strength. D, Communication metrics in infarction versus control within D10, with donor-label permutation nulls. E, Equivalence testing against the ±30-unit margin; the minimum detectable effect at 80% power was 105–213 units. F, Cell-count-matched resampling. G, Donor leave-one-out ranges. Hub metrics are given in Supplementary Table S11, the full matrix in Supplementary Table S12, disease-state metrics and null models in Supplementary Table S13, and region-resolved networks in Supplementary Figure S15. **Alt text:** Seven-panel figure. A, cell-type by cell-type communication network; B, net strength by cell class; C, ranked net strength; D, disease versus control metrics with permutation nulls; E, equivalence testing; F, count-matched resampling; G, donor leave-one-out ranges.

Comparing infarction with control within D10, directional metrics shifted consistently across donor leave-one-out and cell-count-matched resampling, but none exceeded donor-label permutation nulls, and the magnitudes could not be bounded within the pre-specified ±30 arbitrary-unit equivalence margin (minimum detectable effect at 80% power, 105–213 units; Figure 7D–G and Supplementary Table S11). One directional statement survives: the one-sided upper bound on the change in cardiomyocyte output was −1.2 units, excluding any gain in signalling authority beyond the margin, and cardiomyocytes remained net receivers (−69.9 in infarction versus −79.6 in control). The epicardial metric rested on 15 control nuclei and is excluded from inference. We therefore do not interpret these data as a reorganization of cardiac communication; we report an absence of evidence for coupling between identity change and relationship change, not evidence of decoupling. Region-resolved networks and hub metrics are in Supplementary Figure S15 and Supplementary Tables S11–S13.

### The natriuretic secretome is spatially uncoupled from its endothelial receptor

To test whether the stressed cardiomyocyte’s secretome is spatially coherent, we used Visium spatial transcriptomics with cell-type deconvolution (Figure 8A and Supplementary Figure S16; histological-region composition in Supplementary Figure S17). The stressed substate did not activate a broad secretory programme; it specifically up-regulated NPPB (10.7-fold over pooled resting cardiomyocytes) while losing BMP10, a maturation and quiescence factor, and down-regulating VEGFA and DKK3 (Figure 8C). This is targeted, not pleiotropic, secretory rewiring. The cognate receptor NPR1 was expressed most highly by endothelial cells (0.203), not by neighbouring cardiomyocytes or myofibroblasts (Figure 8D).

**Figure 8.**
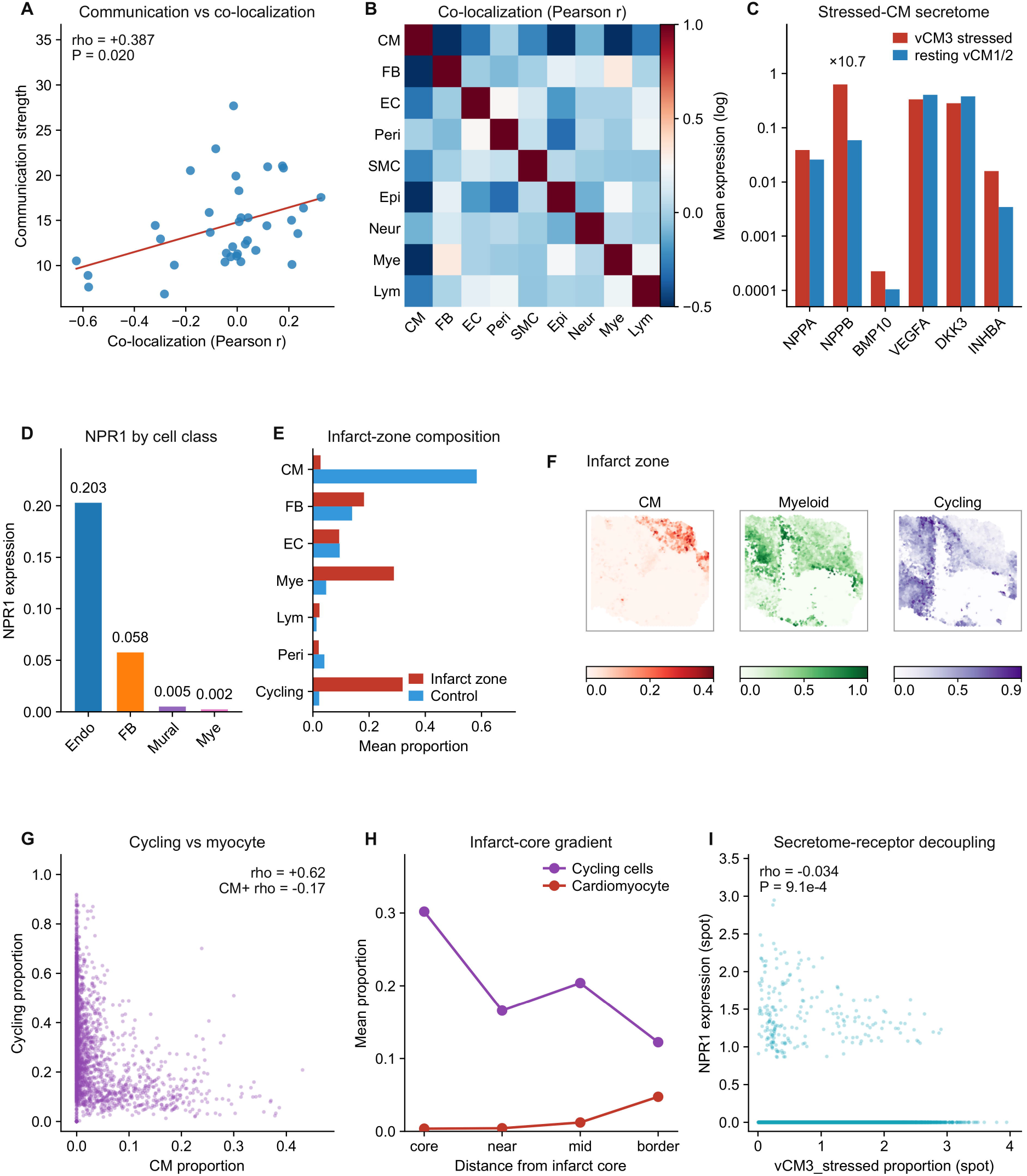
Spatial validation, uncoupling of the natriuretic secretome and infarct-zone reprogramming. A, Communication strength versus spatial co-localization after aligning both matrices to nine major cell classes (Spearman ρ = +0.387, P = 0.020; permutation P = 0.021; donor-slide leave-one-out 0.36–0.39; 36 class pairs). B, Spatial co-localization matrix. C, Secretome of the stressed substate: NPPB induced 10.7-fold, BMP10, VEGFA and DKK3 reduced. D, NPR1 expression by cell class; endothelial cells are the dominant source. E, Infarct-zone composition: cardiomyocyte proportion 0.583 to 0.027, myeloid expansion sixfold, cycling cells fourteenfold. F, Spatial maps of cardiomyocyte, myeloid and cycling-cell proportions in the infarct zone. G, Cycling-cell proportion versus cardiomyocyte content in cardiomyocyte-containing spots (ρ = −0.17, P = 4 × 10□□). H, Infarct-core to border gradient. I, Spot-level relationship between the vCM3_stressed programme and NPR1 expression (ρ = −0.034, P = 9.1 × 10□□); the slope is negligible. Distance-resolved cross-correlations and neighbourhood tests are given in Supplementary Figure S18 and Supplementary Table S15. **Alt text:** Nine-panel figure. A, communication strength versus spatial co-localization; B, co-localization matrix; C, stressed-substate secretome; D, NPR1 expression by cell class; E, infarct-zone composition; F, spatial maps; G, cycling versus cardiomyocyte content; H, infarct-core gradient; I, secretome-receptor decoupling.

NPPB was spatially structured in patches in both the infarct zone (Moran’s I = 0.192, permutation P = 2 × 10□³) and control tissue (I = 0.202, P = 2 × 10□³; Supplementary Figure S18A). Its spot-level abundance tracked cardiomyocyte content far more strongly in the infarct zone (ρ = +0.44) than in control (ρ = +0.09), consistent with cardiomyocytes as the dominant source after infarct-zone collapse. If the ligand engaged its receptor locally, NPPB and NPR1 should be cross-correlated at short range. They were not: the cross-correlation was 0.004 at 0–200 µm and indistinguishable from zero at every distance, whereas COL1A1–ITGB1 showed a strong short-range cross-correlation that decayed with distance (−0.19 at 0–200 µm to +0.01 at 1.6–4 mm; Figure 8B and Supplementary Figure S18B). Spots with the highest NPPB content sat in endothelial-poor neighbourhoods (−3.5 percentage points relative to NPPB-low spots; permutation P = 0.001), an uncoupling that was threefold weaker in control tissue (−1.1 points; Supplementary Figure S18C). A spot-level correlation between the stressed-cardiomyocyte programme and NPR1 was also negative and negligible (ρ = −0.034, P = 9.1 × 10□□; partial ρ = −0.04, P = 0.055 after adjustment for total counts; Figure 8I; spot-level and neighbourhood summaries in Supplementary Figure S18D,E).

The stressed cardiomyocyte therefore loses its identity and redirects secretory output toward a distant endothelial target that is not spatially coupled to it. This is a candidate mechanism, not a demonstrated one: we did not test whether NPPB released by stressed cardiomyocytes engages endothelial NPR1 to produce a functional response. Whether the spatial uncoupling contributes to the clinical dissociation between natriuretic-peptide elevation and functional recovery is a question for functional studies. The present data establish the spatial pattern, and the genetic data establish that the ligand is on a causal path to heart failure.

Spatial analysis also resolved the tissue-level consequence of the identity crisis (Figure 8E). Cardiomyocyte proportion collapsed from 0.583 to 0.027, whereas myeloid cells expanded about sixfold and deconvolution-inferred cycling cells about fourteenfold (Figure 8E,F; Supplementary Table S14; spatial statistics in Supplementary Table S15). At the spot level, cycling-cell proportion correlated positively with cardiomyocyte content in the raw data (ρ = +0.62), but this largely reflected shared tissue density; after restricting to cardiomyocyte-containing spots the association was negative (ρ = −0.17, P = 4 × 10□□; Figure 8G). A spatial gradient from infarct core to border showed cycling cells declining while cardiomyocyte content recovered (Figure 8H). The proliferative response to infarction is therefore a fibro-inflammatory programme, not a regenerative one, which is the tissue-scale expression of the single-cell finding that cardiomyocyte proliferation remains quantitatively minor.

### Erosion is specified to the injured zone and does not generalize to chronic cardiomyopathy

To ask whether identity erosion is a property of injured myocardium or of heart failure generally, we re-analysed an independent cohort of dilated and arrhythmogenic cardiomyopathy (311,418 cardiomyocytes, 79 donors; 18 controls, 52 dilated, 8 arrhythmogenic; [23]; cohort overview in Supplementary Figure S19A–D). In whole cardiomyocytes, IRX4 was not reduced in chronic cardiomyopathy (β = −0.005, 95% confidence interval −0.019 to +0.009; P = 0.48; Figure 9A), and the identity score differed only marginally (β = −0.34, P = 0.041; Figure 9E). By contrast, the stress programme replicated robustly (NPPB β = +0.49, P = 0.007; NPPA P = 0.026; composite stress score β = +0.52, P = 0.008). The sarcomeric arm shifted: MYH6 fell (β = −0.36, P = 0.004) without a change in MYH7, so the MYH6/MYH7 balance moved toward the fetal-like direction (Figure 9B–D,F). The stress and sarcomeric components of the phenotype are therefore general properties of failing myocardium, whereas IRX4 erosion is not.

**Figure 9.**
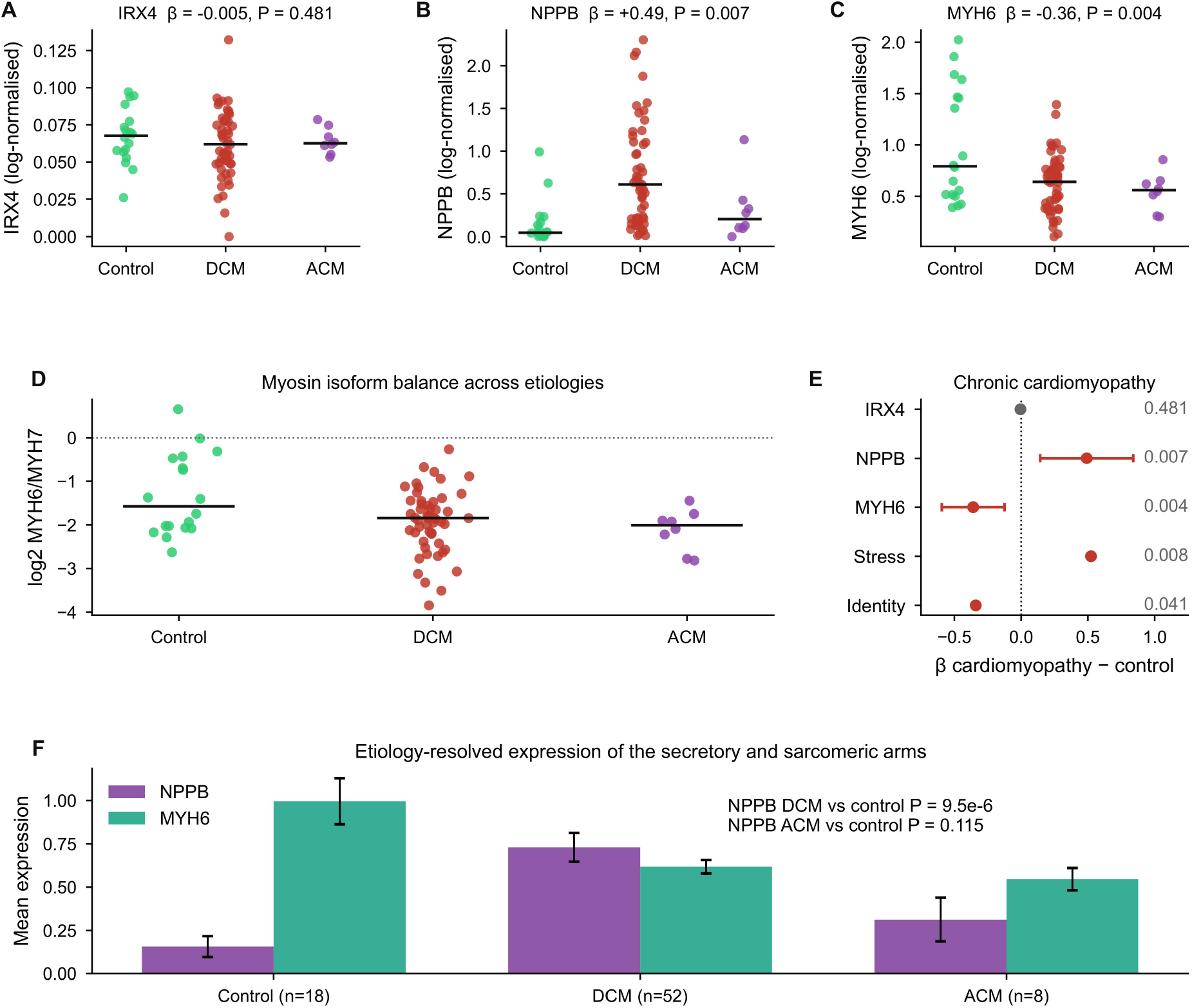
Erosion is specified to the injured zone and does not generalize to chronic cardiomyopathy. A–c, Donor-level IRX4 (A), NPPB (B) and MYH6 (C) expression in an independent cohort of 79 donors (18 controls, 52 dilated cardiomyopathy, 8 arrhythmogenic cardiomyopathy); each point is a donor and bars the median. Disease effects are from ordinary least squares with sex and cell number as covariates. D, Myosin isoform balance (log□ MYH6/MYH7) across aetiologies. E, Disease effects (β with 95% confidence interval) for IRX4, NPPB, MYH6, the stress score and the identity score; red, P < 0.05. F, Aetiology-resolved expression of the secretory (NPPB) and sarcomeric (MYH6) arms. Donor-level statistics are given in Supplementary Table S16, the cohort overview in Supplementary Figure S19A–D, and genetic context in Supplementary Figure S14A–E. **Alt text:** Six-panel figure. A to C, donor-level IRX4, NPPB and MYH6 expression in 79 donors; D, myosin isoform balance; E, disease-effect estimates; F, aetiology-resolved expression.

We interpret this as a boundary condition, not a failed replication. IRX4 erosion is a local, injury-zone event: it is strongest where acute ischaemia coexists with the inflammatory and fibrotic transition, and it is diluted or resolved once the acute niche is no longer present. Chronic cardiomyopathy lacks that niche, and its cardiomyocytes retain IRX4 while activating stress genes. The dissociation is consistent with the spatial data, in which identity loss and NPPB induction co-localize in the ischaemic core, and with the genetic data, in which IRX4 variation does not confer heart-failure risk. The identity crisis is therefore spatially and temporally specified by injury, which is what makes it a candidate target for early intervention, not a generic marker of heart failure.

Together the layers support a two-layer model (Figure 9 and Supplementary Table S16). The state layer is chromatin-encoded identity erosion, confined to the injured zone and accompanied by a targeted natriuretic secretome. The relationship layer, assessed with donor-permutation nulls and equivalence bounds, does not resolve a reorganization at this sample size. The two layers are therefore not coupled in a way this design can detect, and the honest statement is an absence of evidence for coupling, not evidence of decoupling. What survives is a mechanistic claim with three independent supports: transcript, chromatin and genetics for the state; spatial distance for the secretome; and an independent cohort for its boundary.

## Discussion

Infarcted cardiomyocytes undergo a module-wide erosion of ventricular identity that is encoded in chromatin and is unlikely to be a reversible transcriptional fluctuation. Proliferation is not restored: double-positive cycling nuclei remain at 0.011%, and the mitotic response is confined to non-myocytes. Genetic variation separates the identity node from its secretory output, with IRX4 carrying no common-variant association with heart failure and NT-proBNP on a causal path to it. The injured cardiomyocyte adopts a non-regenerative stress identity, and the natriuretic peptide that reports this state is itself a causal signal for adverse outcome.

The mechanistic content of the identity crisis lies in the asymmetry between identity and metabolic programmes. IRX4 is an Iroquois-class homeodomain factor required for ventricular chamber specification[27–29]; its expression is modulated by Nkx2-5 and dHand[29], and its loss in mice causes adult-onset cardiomyopathy preceded by abnormal ventricular gene expression[27]. In the injured human heart, IRX4 activity and locus accessibility fall while ESRRA/ERRα activity rises and fatty-acid oxidation is retained or enhanced. The cell therefore loses the factor that specifies ventricular identity without losing the metabolic programme that defines maturity. This is the opposite of reversion: a fetal cardiomyocyte is glycolytic and proliferative, whereas the injured cardiomyocyte is oxidative and quiescent. The re-expressed fetal genes are downstream stress effectors, not upstream identity determinants, which resolves the paradox that natriuretic-peptide induction has been read as evidence of immaturity.

The genetic dissociation matters for how natriuretic peptides are interpreted. Plasma natriuretic peptides are the most widely used cardiac biomarkers, and their elevation is routinely interpreted as a marker of wall stress. Our cis-pQTL Mendelian randomization indicates that genetically predicted NT-proBNP is on a causal path to heart failure (odds ratio 1.12 per s.d.), which is consistent with a contributory role, not merely a reflective one. At the same time, inherited variation at IRX4 does not confer heart-failure risk, even though IRX4 is the chromatin-defined node of the injured cell. A cell-state marker need not be a disease gene: the state can be induced by injury without common variants at the node altering susceptibility. IRX4 silencing is a recurrent event in other tissues[39], which raises the possibility that the injury-induced loss we describe is a general mechanism of Irx4 repression, not a heart-specific one. This distinction argues against IRX4 as a germline drug target while leaving open the possibility that restoring its activity in the injured zone is therapeutic.

MKI67 detection was enriched 3.7-fold in the ischaemic core, in direction agreement with immunostaining studies[25], yet double-positive nuclei remained at 0.011% and the response was concentrated in a minority of donors. The proliferative response to infarction was dominated by myeloid expansion and by deconvolution-inferred cycling cells that carry no lineage information and sit in cardiomyocyte-depleted regions. The heart’s response to infarction is therefore a fibro-inflammatory programme, and any regenerative strategy must either convert that programme or act downstream of it[40]. The identity crisis offers a specific lever: because the state is chromatin-encoded and proliferatively silent, inducing proliferation without addressing it may produce cycling cells that retain an eroded identity.

NPPB was induced while BMP10, a maturation and quiescence factor, was lost, and the ligand was spatially uncoupled from its endothelial receptor. If this uncoupling holds functionally, it would place the natriuretic signal in a paracrine void: the peptide is released but its endothelial target is not adjacent. This would explain why natriuretic-peptide elevation tracks adverse outcome without producing a compensatory response, and it suggests that the NPPB–NPR1 axis should be tested as a target, not assumed to be a readout. The claim is a spatial one; functional engagement remains to be demonstrated.

Several limitations apply. All human data are re-analyses of public datasets, and the infarct cohort lacks time-since-infarction annotation, so acute and chronic stages cannot be resolved; region-based stratification is the only temporal proxy. The maturation reference draws on single-cell descriptions of human cardiac maturation[41]; the axis samples mid-gestation (19–20 weeks), although first-trimester hearts showed the same ventricular MYH7 dominance. The transcript-level IRX4 difference is modest (ratio 0.84, donor-paired P = 0.067), so the case rests on convergence across activity, chromatin, spatial gradient and genetic context, and not on expression magnitude; protein-level measurement would strengthen it. The chromatin analysis uses gene-activity scores, not peak-level counts (Supplementary Figure S20A–D), so motif-level inference was not possible, and the control zone comprises four donors, placing the donor-level test near its resolution floor. Communication inference is correlational and underpowered, with a minimum detectable effect of 105–213 units; the ligand–receptor compendium carries no cell-type-specific weights or diffusion constraints and therefore captures potential, not realized, signalling. The Mendelian randomization rests on a single cis instrument, so directional pleiotropy cannot be tested formally; the estimate was robust to the conditional pQTL effect but not to violations of the exclusion restriction. The spatial analysis rests on one Visium section per group, and the cross-cohort comparison cannot separate chronicity from aetiology. No gain- or loss-of-function perturbation of IRX4 was performed, so its causal role rests on convergence across layers, not on direct intervention. The study is observational and does not establish a therapeutic effect.

Together, these layers define an injured cardiomyocyte state that is chromatin-encoded, proliferatively silent and spatially confined to the ischaemic zone, and they separate the identity node from the natriuretic output that reports it. The state is a candidate target for early intervention, but the present evidence is observational and genetic; direct perturbation of IRX4 in human myocardium remains the decisive test.

## Materials and Methods

### Data sources

All human data are public and were downloaded from CELLxGENE (cellxgene.cziscience.com). The adult human heart reference (486,134 cells; Litviňuková et al.[20]) was used as the communication baseline (D1). Human adult and first-trimester fetal hearts (60,668 cells)[42] (D2) and a left-ventricular maturation axis spanning 19–20 gestational weeks (D5) provided the developmental reference; a mouse perinatal series (49,769 cells; D4) provided the cross-species comparison. The Human Cell Atlas cardiac-niche collection (704,296 cells; Kanemaru et al.[21]) supplied the eight-region adult reference and the cardiomyocyte substates (D7). The infarcted human heart multi-omic dataset (Kuppe et al.[22]) supplied single-nucleus RNA-seq (191,795 nuclei), single-nucleus ATAC-seq (46,086 nuclei) and Visium sections (D10). An independent cardiomyopathy cohort (Reichart et al.[23]; 311,418 cardiomyocytes, 79 donors) was used for cross-aetiology comparison. Genetic data were FinnGen release 13 summary statistics for eight phenotypes[30] and the deCODE plasma proteome quantitative trait loci[33]. Open Targets release 26.09 provided gene–disease association scores[43].

### Cell-type harmonisation and cardiomyocyte extraction

Raw annotations were harmonised into ten consensus classes (cardiomyocyte, fibroblast, endothelial, smooth muscle, pericyte, myeloid, lymphoid, neuronal, epicardial, adipocyte) by keyword mapping. Cardiomyocytes were extracted for maturation analysis (38,295 human, 17,047 mouse). Gene symbols were unified to uppercase for cross-species comparison.

### Maturation trajectory

Five programmes were scored as the mean expression of curated gene sets (proliferation: MKI67, TOP2A, CCNB1, PCNA, CCNA2; dedifferentiation: NPPA, NPPB, ANKRD1, MYH6, MYH7; glycolysis: HK2, PKM, LDHA, PDK1, SLC2A1; fatty-acid oxidation: CPT1B, PDK4, ACADM, CD36; maturation: MYL2, TNNI3, RYR2, CKM, ATP2A2). Cells were ordered by developmental maturity, and monotonicity was assessed by Spearman correlation across stage means.

### IRX4 pseudobulk quantification

Raw counts were taken from the CELLxGENE objects and summarised per donor and per state. For each donor and population, IRX4 counts were summed and divided by the summed total counts of the same cells to give copies per million; the detection rate was the fraction of nuclei with at least one IRX4 transcript. Ratios were bootstrapped over donors (2,000 resamples).

### Transcription-factor activity

Activity was inferred with decoupler v2.2.0 using the CollecTRI human regulon (1,185 factors) and univariate linear models, comparing fetal, adult and infarcted cardiomyocytes. Fetal switches were defined as log□ fold change > 0.5 with false-discovery rate < 0.05 (fetal versus adult). The focal factors were re-inferred with AUCell and multivariate linear models as a method-independence check.

### Chromatin accessibility and donor-aware models

Single-nucleus ATAC-seq gene-activity scores were computed from the CELLxGENE object (46,086 nuclei). Cardiomyocyte-lineage nuclei (cardiac muscle myoblast; 21,923) were assigned to control, border (myogenic), fibrotic and ischaemic zones; nuclei from the control region of infarcted donors were assigned to the control zone. Per-gene activity was z-scored across nuclei for module scores. Zone effects were estimated by ordinary least squares with donor-clustered robust standard errors (20 donors), and stability was assessed by leave-one-donor-out resampling. Regulon enrichment was computed as the mean target-gene Δlog□ fold change between ischaemic and control zones for 750 CollecTRI factors with at least five measurable targets, with Benjamini–Hochberg correction.

### Genetic analysis

FinnGen release 13 summary statistics were streamed and the IRX4 interval (chr5:1,850,000–1,950,000, GRCh38) extracted for eight phenotypes. Variants with minor-allele frequency below 1%, standard error above 0.5 or missing effect estimates were excluded, leaving 606–608 variants per phenotype. No variant reached P < 5 × 10□□. The lead conduction variant rs260399 was analysed directly. The chronic obstructive pulmonary disease summary-statistics file was truncated in the source download; because the two intervals analysed here lie on chromosomes 1 and 5, which precede the truncation point, the variants used for that phenotype were complete.

For Mendelian randomization, rs198389 was used as a cis instrument for NT-proBNP (deCODE; β = 0.259 s.d. per G allele, P = 5.7 × 10□²¹□; minor-allele frequency 0.40)[33]. Alleles were harmonised on the ALT allele, which was the effect allele in both datasets (FinnGen ALT frequency 0.364–0.366). The Wald ratio was computed as β_out/β_exp with a first-order standard error, and the instrument strength as F = (β_exp/se_exp)², where se_exp was derived from the reported P value; variance explained was approximated as 2f(1−f)β². A sensitivity analysis repeated the estimate with the conditional pQTL effect (β = 0.090). Because a single instrument was available, heterogeneity and directional-pleiotropy tests could not be performed; the estimate is therefore reported as a cis-MR estimate under the standard instrumental-variable assumptions[34,35].

### Communication analysis

Ligand–receptor pairs were obtained from OmniPath (ligrecextra; 6,657 interactions), mapped to gene symbols through the Human Protein Atlas (4,366 mapped pairs; 4,349 expressed in D1). Communication was quantified across the 447,885 D1 cells that expressed at least one ligand or receptor gene. For each expressed pair, strength between sender and receiver classes was the product of the mean ligand expression in the sender and the mean receptor expression in the receiver, summed over pairs. Disease comparisons were performed within D10 with donor-label permutation nulls, cell-count-matched resampling, donor leave-one-out and equivalence testing (margin ±30 units).

### Spatial analysis

Visium spots were deconvolved into cell-type proportions (provided with D10) and aggregated into nine classes. Spatial autocorrelation of NPPB was quantified by Moran’s I on a six-nearest-neighbour graph with 499 permutations. Ligand–receptor spatial coupling was quantified as the Pearson cross-correlation between ligand abundance at spot i and receptor abundance at spot j, averaged over both directions, in distance bins from 0–200 µm to 1.6–4 mm; COL1A1–ITGB1 served as a positive control. Endothelial content in the six nearest neighbours of NPPB-high (upper quartile) versus NPPB-low spots was compared by permutation (999 shuffles). Co-localization matrices and spot-level partial correlations controlled for total counts.

### Cross-cohort analysis

Donor-level pseudobulk values for 79 donors were computed from the cardiomyopathy cohort (Reichart et al.[23]) as mean log-normalised expression across cardiomyocytes. Disease effects were estimated by ordinary least squares with sex and cell number as covariates; aetiology-specific comparisons used Mann–Whitney tests against controls. Stress and identity scores were the mean of z-scored constituent genes across donors.

### Statistics and reproducibility

Monotonicity and correlation were assessed with Spearman rank correlation; group comparisons with Mann–Whitney U or Wilcoxon tests; multiple testing was controlled by Benjamini–Hochberg false discovery rate. Donor-level summaries (mean and 95% confidence interval) are displayed and donor-aware tests are used by default: donor-clustered robust standard errors for cross-dataset fits and donor-label permutation for network and spatial comparisons. Cell- and spot-level P values describe within-dataset variation and are interpreted as exploratory. No statistical method was used to predetermine sample size; all public datasets passing the original authors’ quality control were analysed. Replicate numbers refer to independent donors or sections and are stated in each legend.

## Data Availability

The datasets analysed here are publicly available on CELLxGENE (https://cellxgene.cziscience.com); dataset identifiers D1–D10 and the corresponding original publications are listed in Supplementary Table S1. FinnGen release 13 summary statistics are available at https://r13.finngen.fi/, and the deCODE plasma proteome pQTLs in the supplementary tables of ref. [33]. Open Targets release 26.09 is available at https://platform.opentargets.org/. All derived tables underlying the figures are provided as Supplementary Tables S1–S16.

The analysis pipeline (dataset retrieval, preprocessing, and figure- and table-generating scripts) is provided as Supplementary file Analysis_Code_v38.zip and is archived at Zenodo under the concept DOI 10.5281/zenodo.24615494; the version accompanying this manuscript will be deposited under the same concept DOI before publication.

## CRediT author statement

**Shigang Qiao:** Conceptualization, Writing – original draft, Writing – review & editing, Visualization. **Lingmei Li:** Methodology, Software, Formal analysis, Data curation, Writing – review & editing, Visualization. **Chunyan He:** Conceptualization, Writing – review & editing, Supervision. **Wei Li:** Conceptualization, Methodology, Software, Formal analysis, Writing – review & editing, Visualization, Supervision, Funding acquisition. All authors have read and approved the final manuscript.

## Ethics Statement

Not applicable. This study analysed only publicly available, de-identified human datasets; no human participants, human tissue or animals were involved.

## Acknowledgments

We thank the investigators of the CELLxGENE, FinnGen, deCODE and Open Targets resources, and the participants and donors whose samples underlie the public datasets analysed here.

## Abbreviations

AUCell: area under the curve for regulon activity
CI: confidence interval
CPM: copies per million
FDR: false discovery rate
IZ: infarct zone
MLM: multivariate linear model
MR: Mendelian randomization
OR: odds ratio
pQTL: protein quantitative trait locus
snATAC: single-nucleus assay for transposase-accessible chromatin
TF: transcription factor
ULM: univariate linear model.

## Supplementary Material

Supplementary material is available online and is cited in the text.

Supplementary Figure S1. Dataset overview. Cell counts, modalities and cell-type composition of the six integrated datasets.

Supplementary Figure S2. Maturation-trajectory support. Programme-gene heat map and mouse trajectory.

Supplementary Figure S3. Cardiomyocyte substate composition. Cell counts across the twelve cardiomyocyte substates.

Supplementary Figure S4. Identity-gene expression across substates. Mean expression of eight identity and stress genes across the twelve substates.

Supplementary Figure S5. Rare-event proliferation counts. MKI67 detection by zone and donor, and double-positive nuclei.

Supplementary Figure S6. Fetal transcription-factor programmes are not re-engaged after infarction. Volcano plot and fetal switch factors.

Supplementary Figure S7. Transcription-factor activity heat map. Top differentially active factors between fetal and adult cardiomyocytes.

Supplementary Figure S8. Transcription-factor activity cross-validation. ULM, AUCell and MLM estimates for IRX4, ESRRA and TEAD1.

Supplementary Figure S9. Per-nucleus sequencing depth confounds zone-level transcript detection. Depth and detection by zone.

Supplementary Figure S10. Chromatin regulon enrichment across 750 CollecTRI regulons. Distribution of mean target log□ fold change between ischaemic and control zones.

Supplementary Figure S11. MYC-target and IRX4-locus accessibility co-variation. Spearman ρ by zone and within-donor values.

Supplementary Figure S12. Donor-level chromatin accessibility. Donor × zone IRX4 gene activity, the paired contrast (not estimable; no donor contributed both zones) and sampling depth.

Supplementary Figure S13. Variant quality control at the IRX4 locus. Variants before and after quality control.

Supplementary Figure S14. Genetic context of the IRX4 locus. Regional association plots for additional phenotypes and Open Targets gene–disease profiles.

Supplementary Figure S15. Region-specific communication networks. Ligand–receptor networks resolved by anatomical region.

Supplementary Figure S16. Deconvolved composition of infarct-zone and control Visium sections. Proportion of spots assigned to each cell class.

Supplementary Figure S17. Histological-region cell-type composition. Deconvolved proportions across histological regions of the D7 Visium atlas.

Supplementary Figure S18. Spatial statistics of the natriuretic secretome. Moran’s I, distance-resolved cross-correlation and neighbourhood composition.

Supplementary Figure S19. The independent cardiomyopathy cohort. Donor-level IRX4, NPPB and stress scores by disease group, and the fraction of vCM3_stressed-like cardiomyocytes.

Supplementary Figure S20. Zone stratification of the chromatin and communication layers. Accessibility and network metrics by zone.

Supplementary Table S1. Inventory of the public datasets and genetic resources used in this study.

Supplementary Table S2. Programme scores versus developmental maturity.

Supplementary Table S3. Marker expression, identity–stress scores and IRX4 pseudobulk across cardiomyocyte substates.

Supplementary Table S4. Programme scores and donor-aware statistics in fetal, adult and infarcted cardiomyocytes; rare-event counts.

Supplementary Table S5. Transcription-factor activity (decoupler, CollecTRI) across states and inference algorithms.

Supplementary Table S6. Differential transcription-factor activity between fetal and adult cardiomyocytes.

Supplementary Table S7. Single-nucleus ATAC gene-activity values, donor means and donor-clustered tests.

Supplementary Table S8. Donor-level chromatin accessibility statistics and regulon enrichment.

Supplementary Table S9. IRX4 locus scan in FinnGen release 13: lead variants, rs260399 effects and the quality-controlled heart-failure interval.

Supplementary Table S10. cis-pQTL Mendelian randomization: instrument, harmonisation and outcome-specific estimates.

Supplementary Table S11. Communication hub metrics for the D1 structural baseline.

Supplementary Table S12. Cell-type by cell-type net communication strength in D1.

Supplementary Table S13. D10 communication metrics, equivalence tests, null models and leave-one-out ranges.

Supplementary Table S14. Deconvolved cell-type composition of the infarct-zone and control Visium sections.

Supplementary Table S15. Spatial statistics for NPPB and the endothelial niche: Moran’s I, distance-resolved cross-correlation and neighbourhood composition.

Supplementary Table S16. Donor-level cardiomyopathy cohort statistics and aetiology-resolved expression.

Key Resources Table. Datasets, software, plasmids and reagents used in this study.

