## Supplementary material for "Chromatin-encoded IRX4 identity erosion bounds cardiomyocyte renewal in the infarcted human heart": Supplementary_Material_v38.pdf

#### Contents


- Supplementary Figures S1–S20 (legend on each page)
- Supplementary Tables S1–S16 (provided as Supplementary_Tables_S1-S16.zip)
- Key Resources Table


This file contains the supplementary figures with their legends and an index of the supplementary tables. Supplementary tables are provided as a separate archive of editable worksheets.

#### Supplementary Figure S1

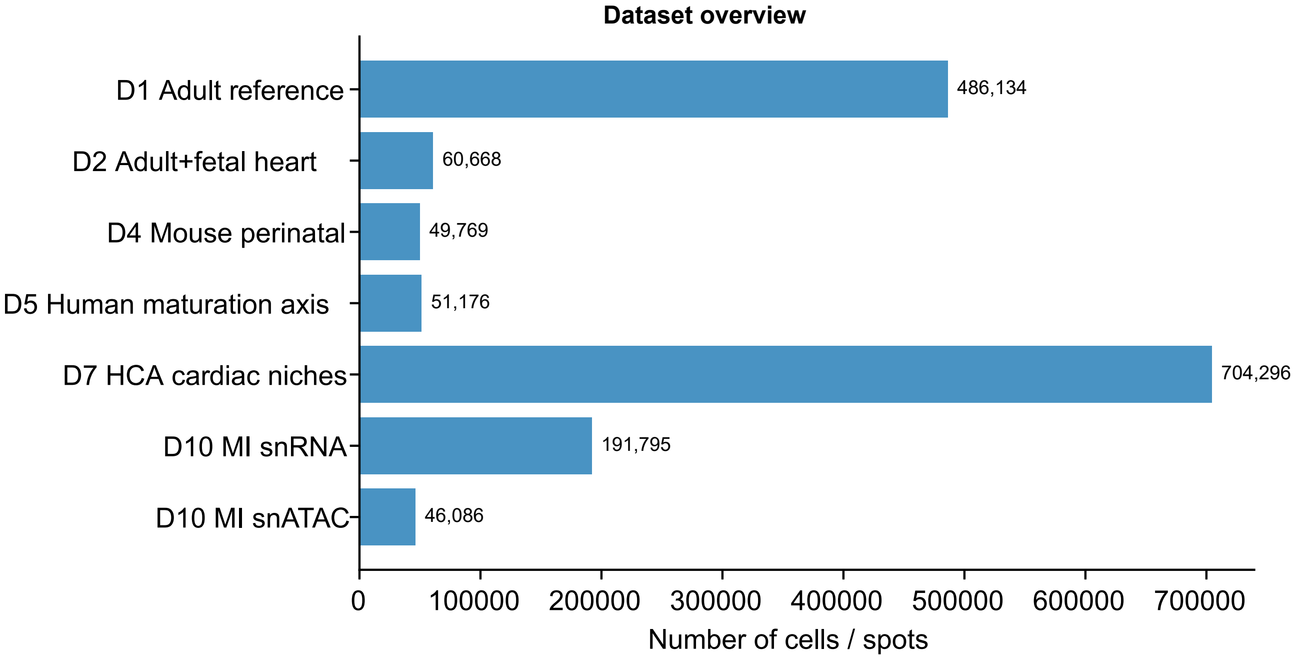


Figure S1. Dataset overview. Cell counts, modalities and cell-type composition of the six integrated datasets. Bars give the number of cells or nuclei passing quality control.

A

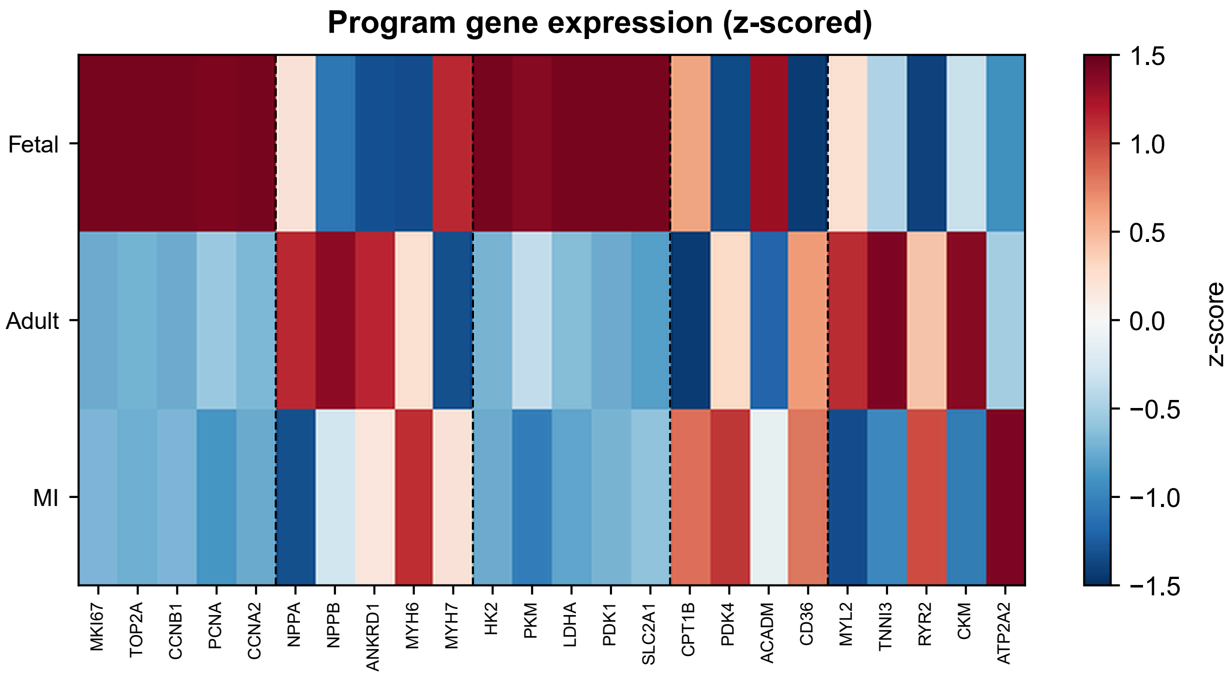


B

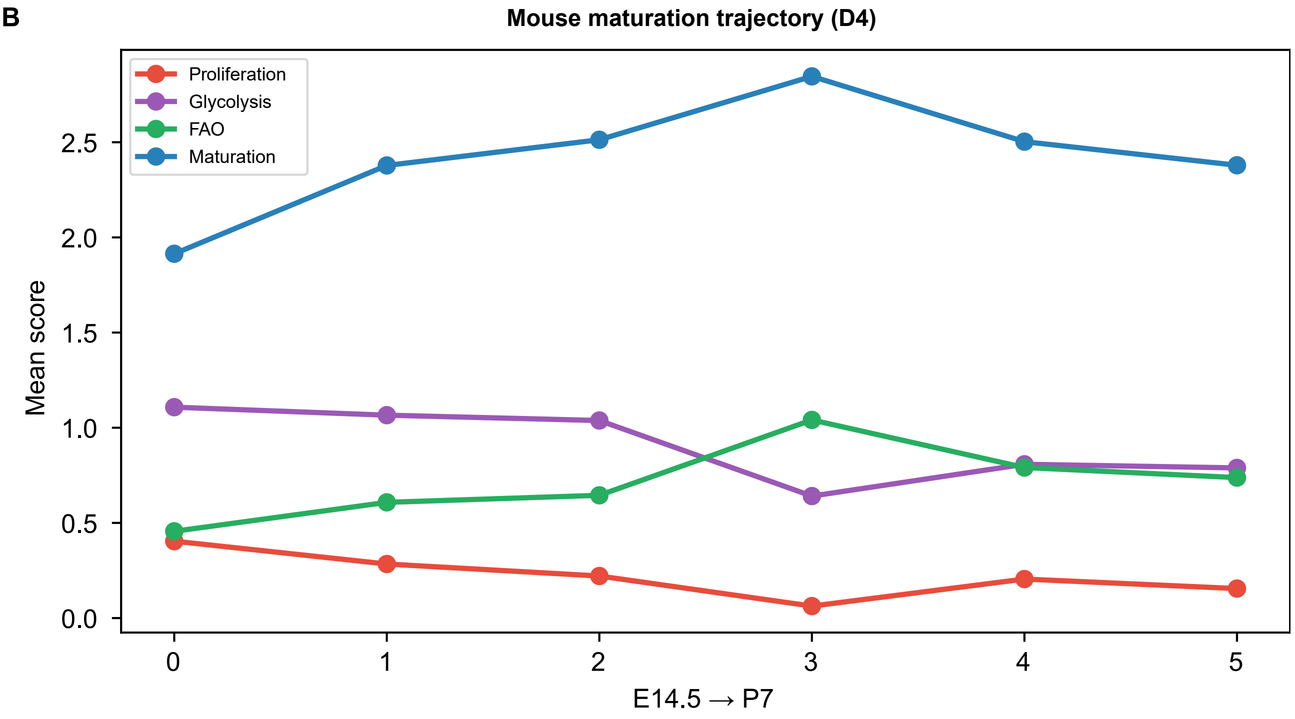


Figure S2. Maturation-trajectory support. (A) Programme-gene heat map across human maturity stages. (B) Mouse perinatal trajectory.

**Supplementary Figure S3**

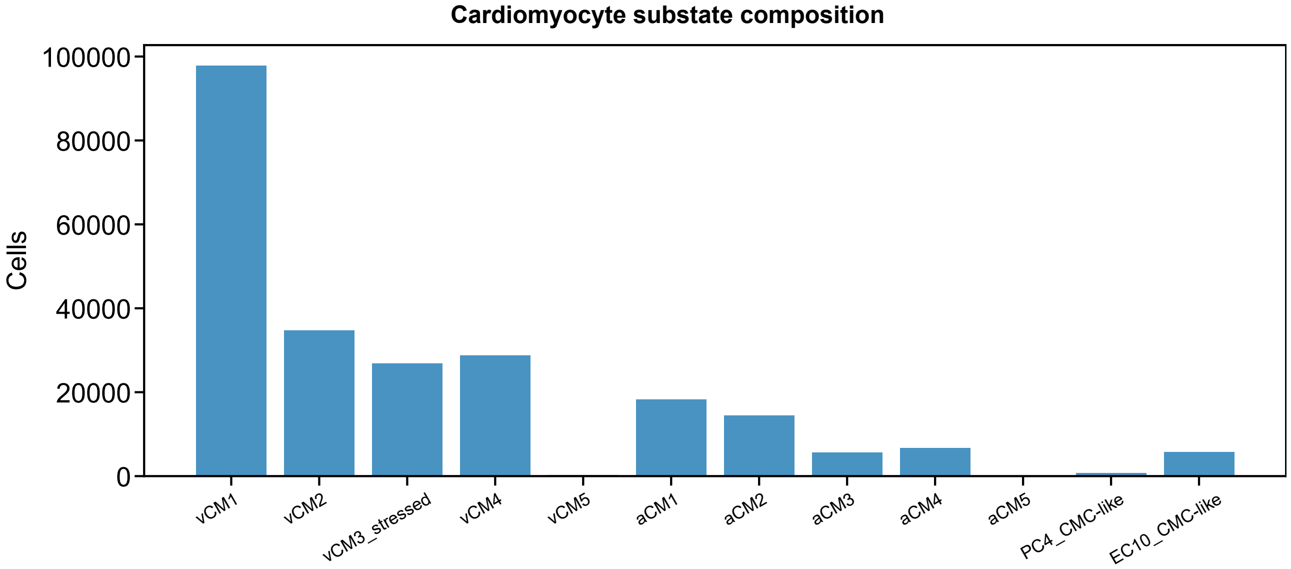


Figure S3. Cardiomyocyte substate composition. Cell counts of the twelve cardiomyocyte substates, including vCM3_stressed.

#### Supplementary Figure S4

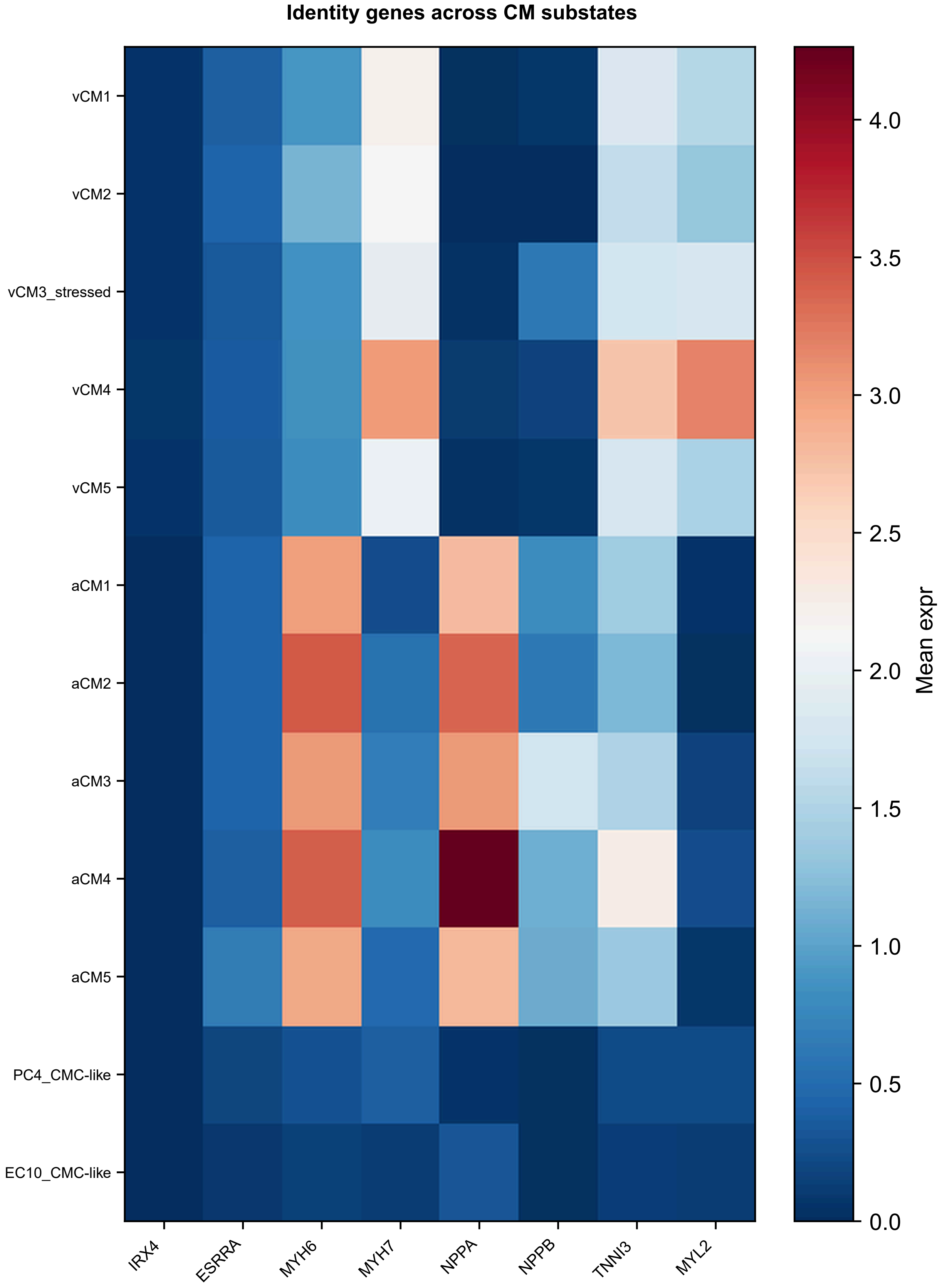


Figure S4. Identity-gene expression across substates. Mean expression of IRX4, ESRRA, MYH6, MYH7, NPPA, NPPB, TNNI3 and MYL2 across the twelve substates.

**Supplementary Figure S5**

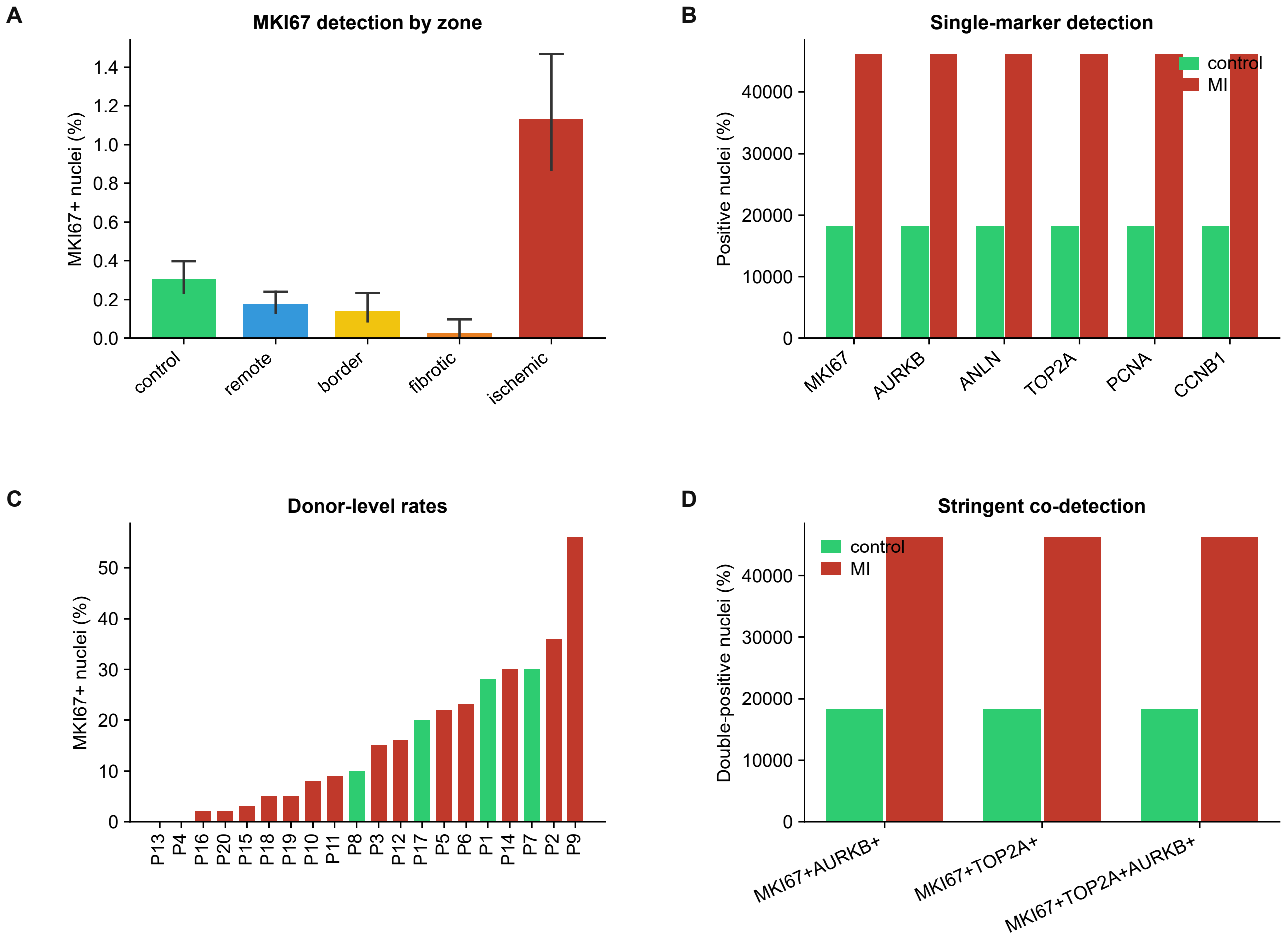


Figure S5. Rare-event proliferation counts. (A) MKI67 detection by zone with Wilson 95% confidence intervals. (B) Donor-level counts. (C) Double-positive nuclei. (D) Zone-resolved comparison.

**Supplementary Figure S6**

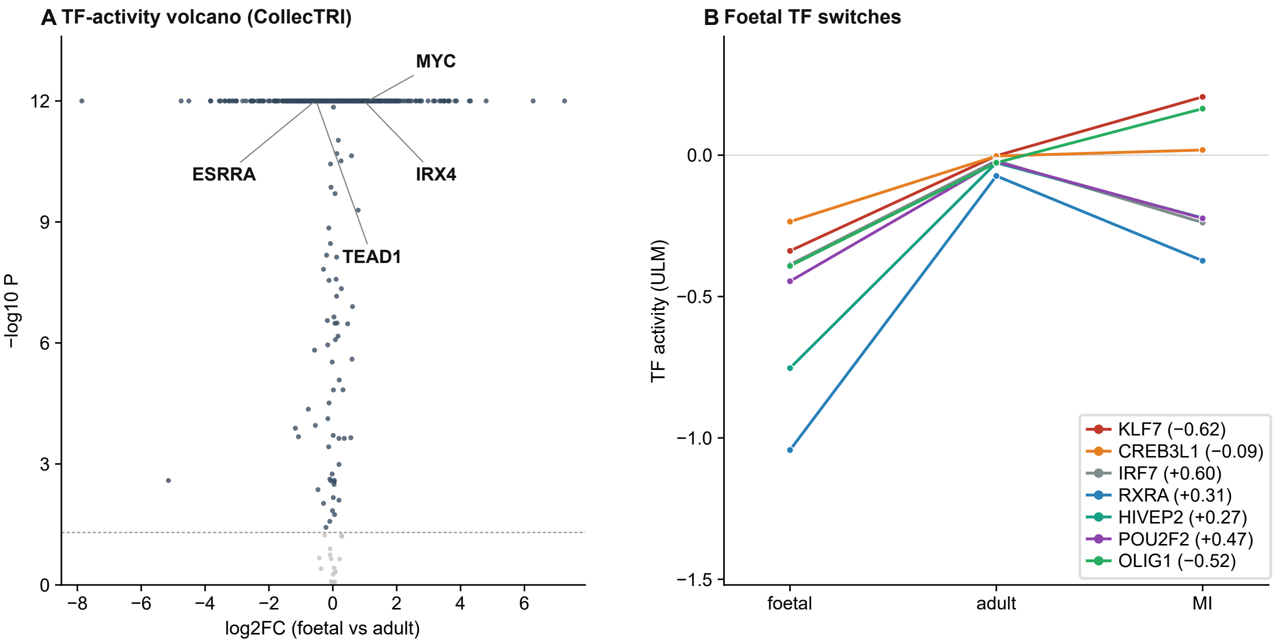


Figure S6. Fetal transcription-factor programmes are not re-engaged after infarction. (A) Transcription-factor activity volcano (CollecTRI). (B) Fetal switch factors in infarcted cardiomyocytes.

#### Supplementary Figure S7

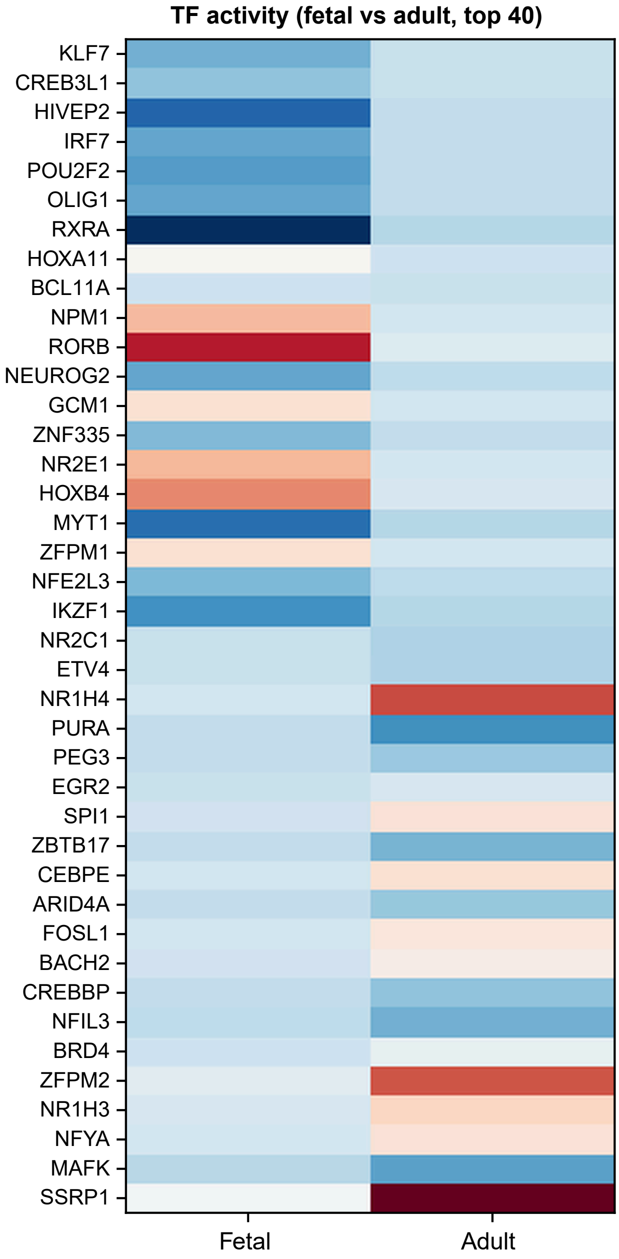


Figure S7. Transcription-factor activity heat map. Top differentially active factors between fetal and adult cardiomyocytes.

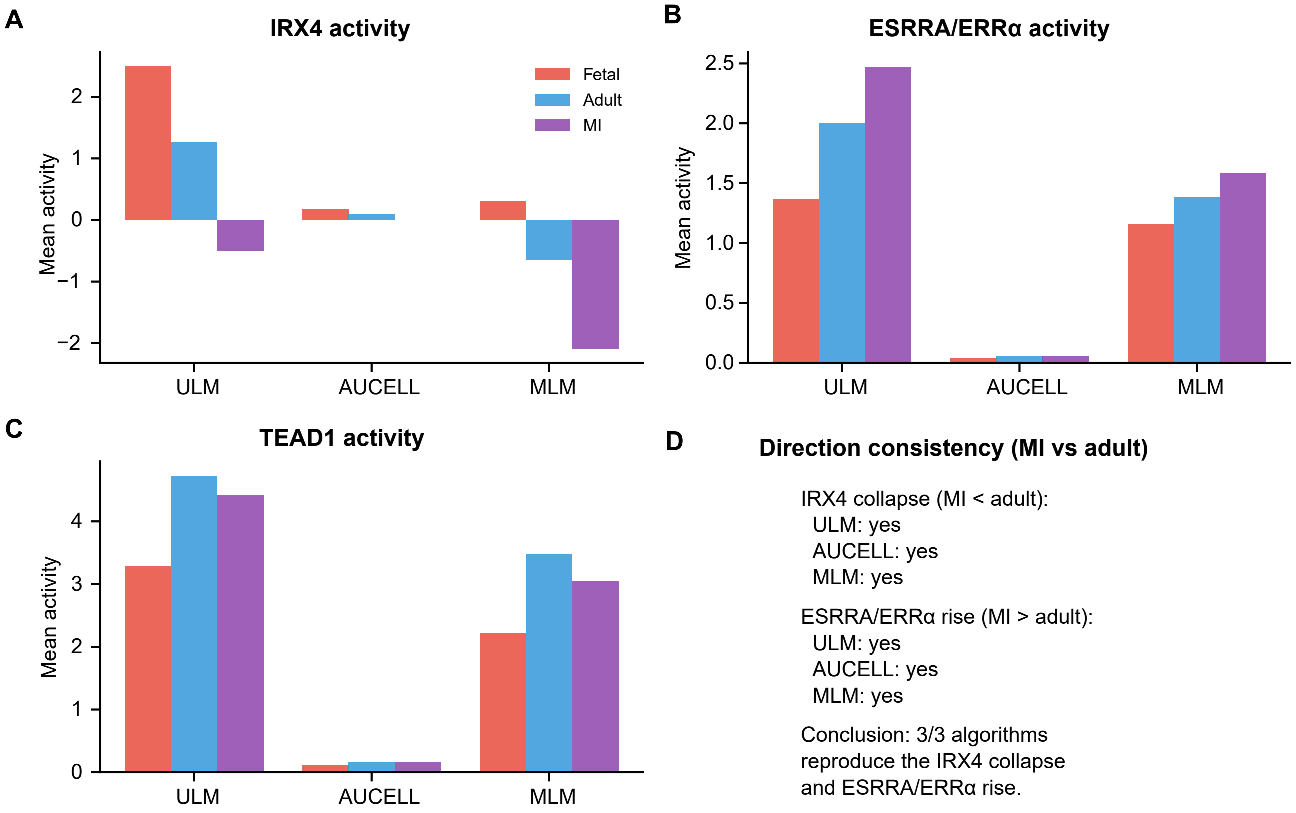


Figure S8. Transcription-factor activity cross-validation. (A) IRX4, (B) ESRRA/ERRα and (C) TEAD1 activity estimated by univariate linear models (ULM), AUCell and multivariate linear models (MLM) in fetal, adult and infarcted cardiomyocytes. (D) Direction consistency of the IRX4 collapse and ESRRA/ERRα rise across the three algorithms.

### Supplementary Figure S9

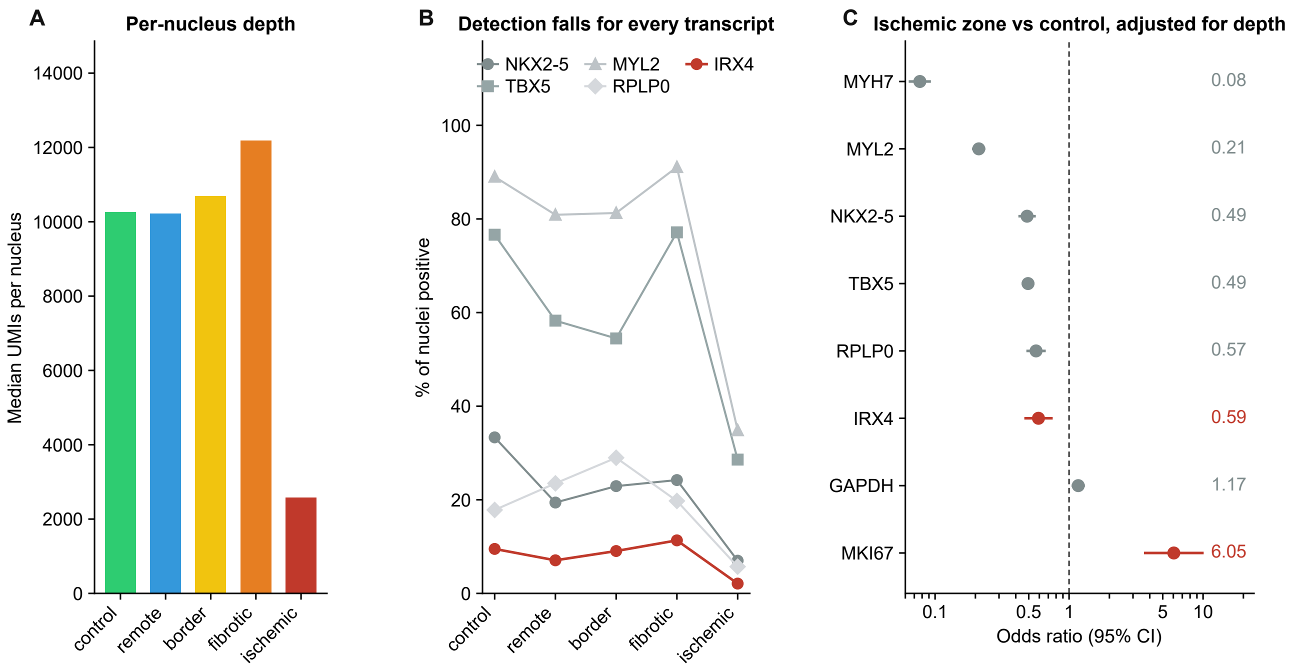


Figure S9. Per-nucleus sequencing depth confounds zone-level transcript detection. (A) Median unique molecular identifiers per nucleus by zone. (B) Detection rate of identity transcripts. (C) Depth-adjusted odds ratios.

**Supplementary Figure S10**

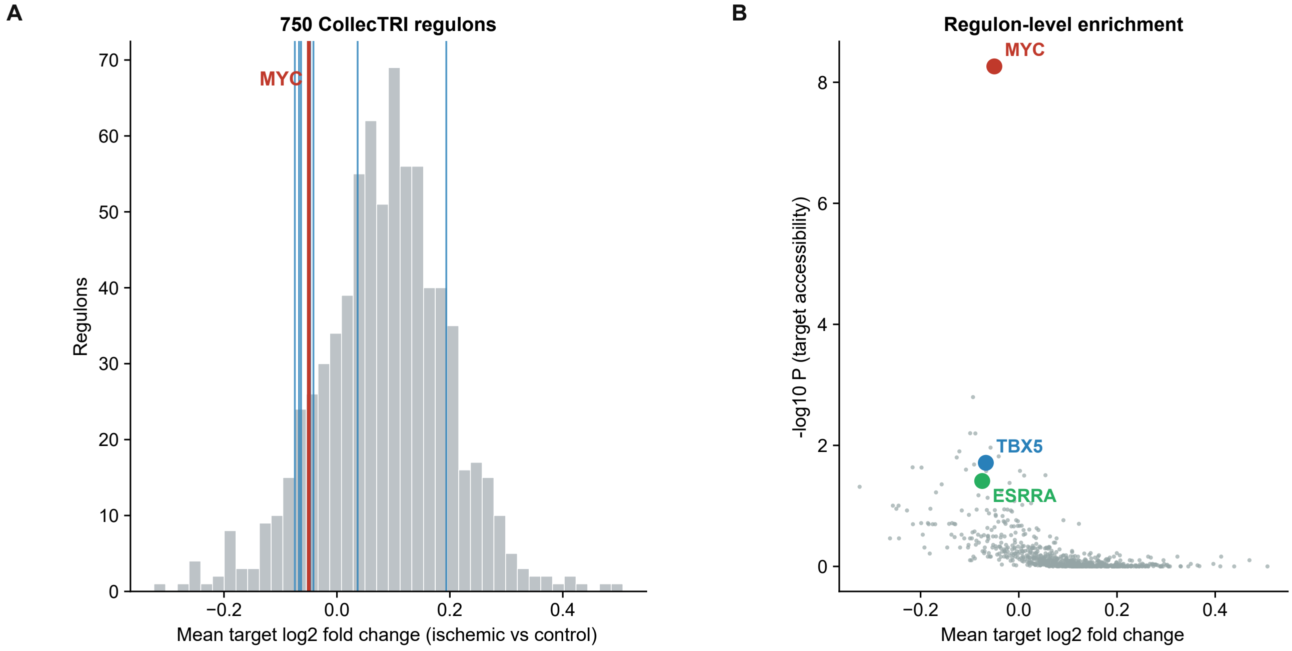


Figure S10. Chromatin regulon enrichment across 750 CollecTRI regulons. (A) Distribution of mean target $\log_2$ fold change between ischaemic and control zones across 750 regulons; MYC is the only factor passing correction. (B) Regulon-level enrichment (target accessibility) versus mean target $\log_2$ fold change, with MYC, TBX5 and ESRRA highlighted.

#### Supplementary Figure S11

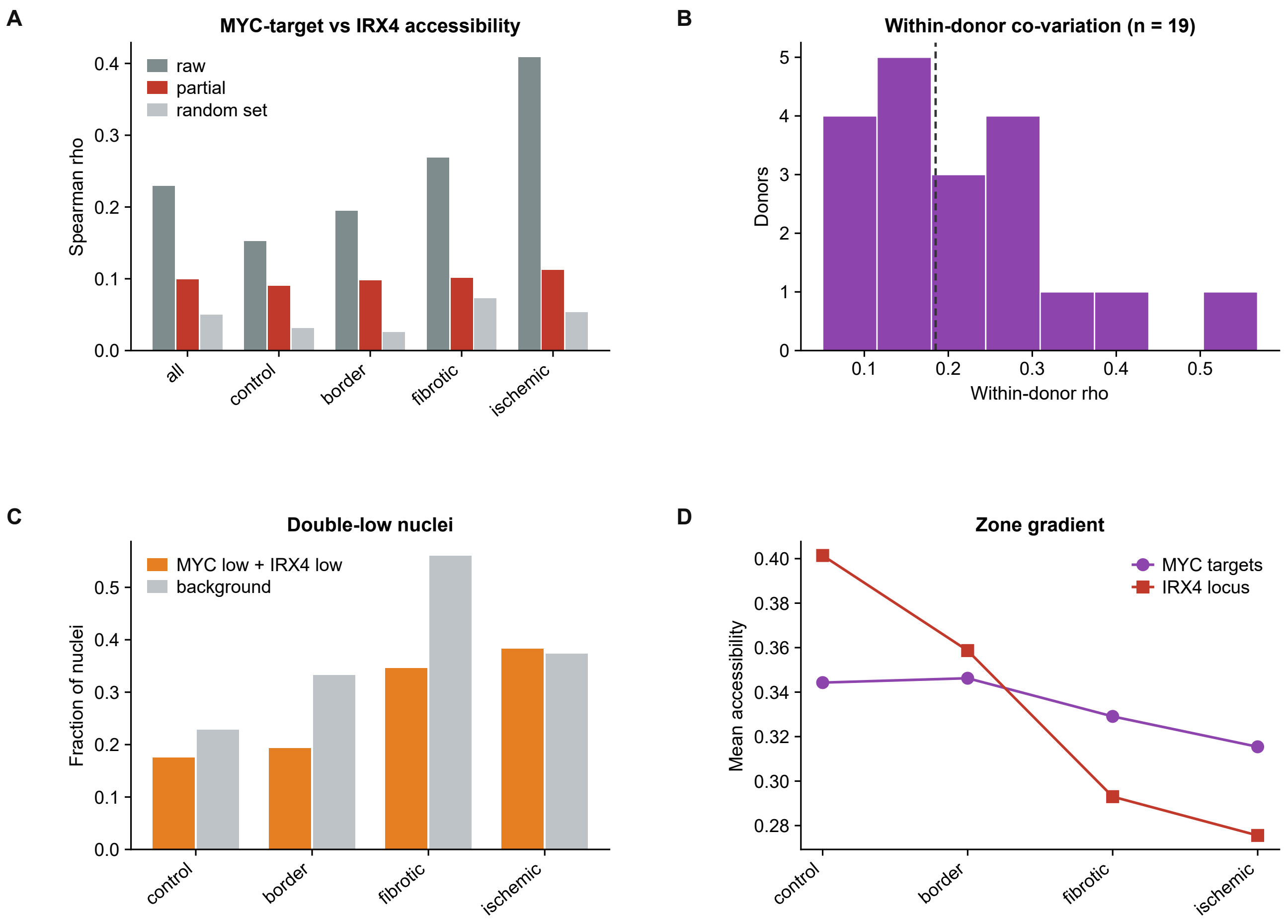


Figure S11. MYC-target and IRX4-locus accessibility co-variation. (A) Spearman correlation between MYC-target and IRX4-locus accessibility across nuclei, before (raw) and after partialling out total counts (partial), compared with random gene sets in each zone. (B) Within-donor correlation coefficients (n = 19 donors). (C) Fraction of nuclei with low MYC-target and low IRX4 accessibility (double-low) versus background. (D) Zone gradient of mean accessibility for MYC targets and the IRX4 locus.

### Supplementary Figure S12

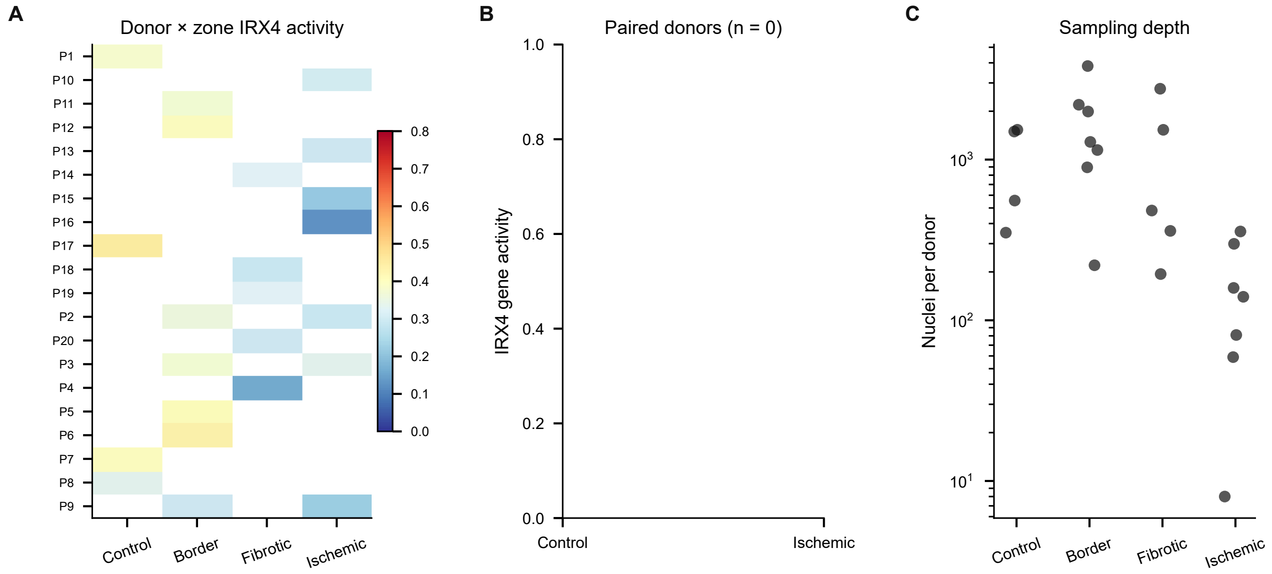


Figure S12. Donor-level chromatin accessibility. (A) Donor × zone IRX4 gene activity. (B) Paired control versus ischaemic values; no donor contributed both a control and an ischaemic sample (n = 0 pairs), so the paired contrast is not estimable. (C) Sampling depth.

**Supplementary Figure S13**

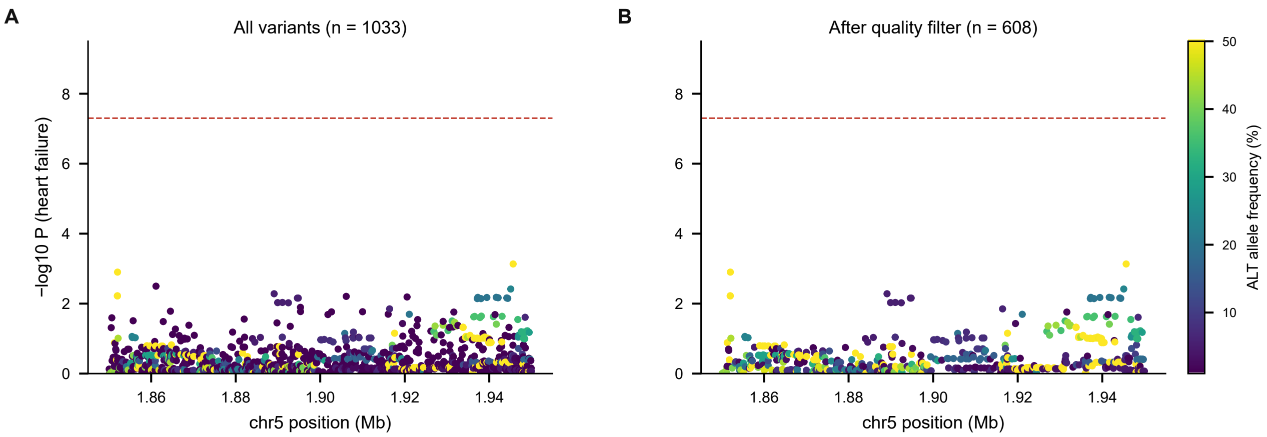


Figure S13. Variant quality control at the IRX4 locus. (A) All variants in the interval. (B) After exclusion of variants with minor-allele frequency below 1%, standard error above 0.5 or missing effect estimates. Colour gives ALT allele frequency.

#### Supplementary Figure S14

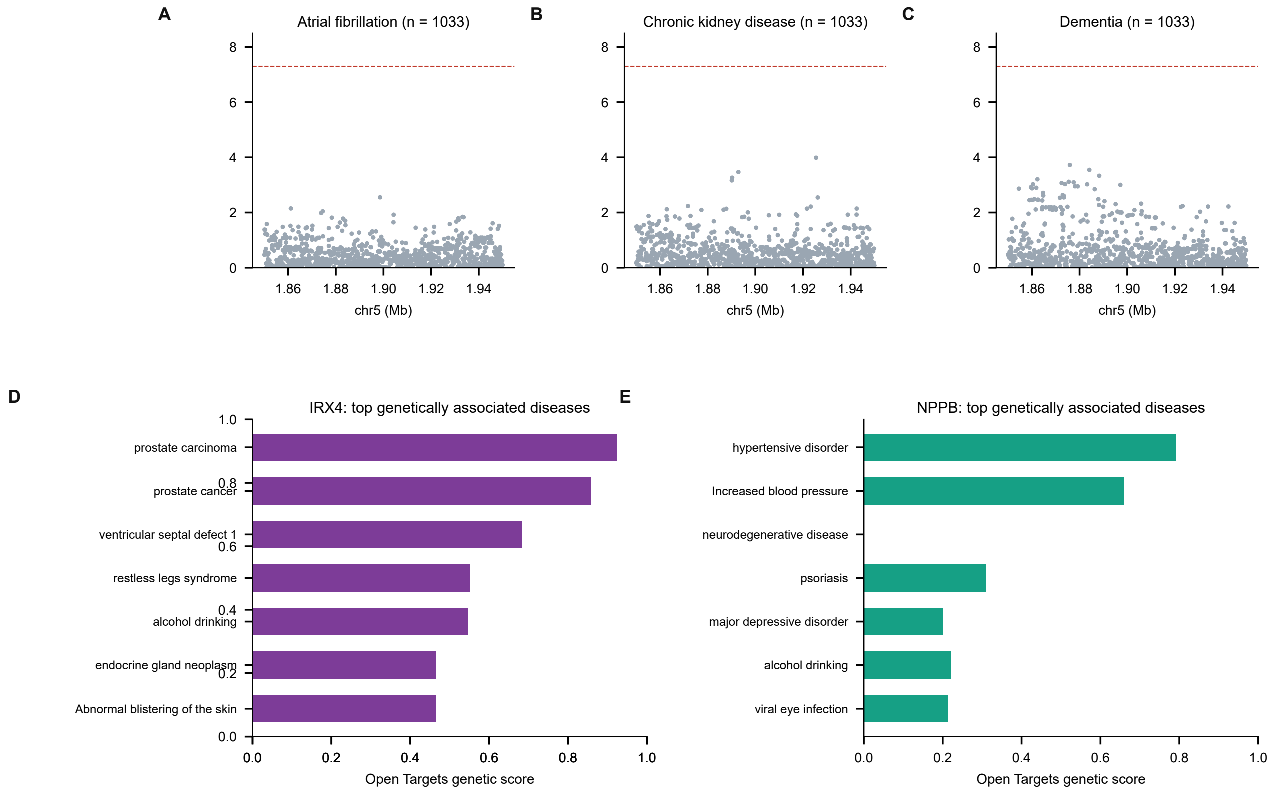


Figure S14. Genetic context of the IRX4 locus. (A–C) Regional association plots for atrial fibrillation, chronic kidney disease and dementia. (D,E) Open Targets 26.09 genetic-association profiles for IRX4 and NPPB.

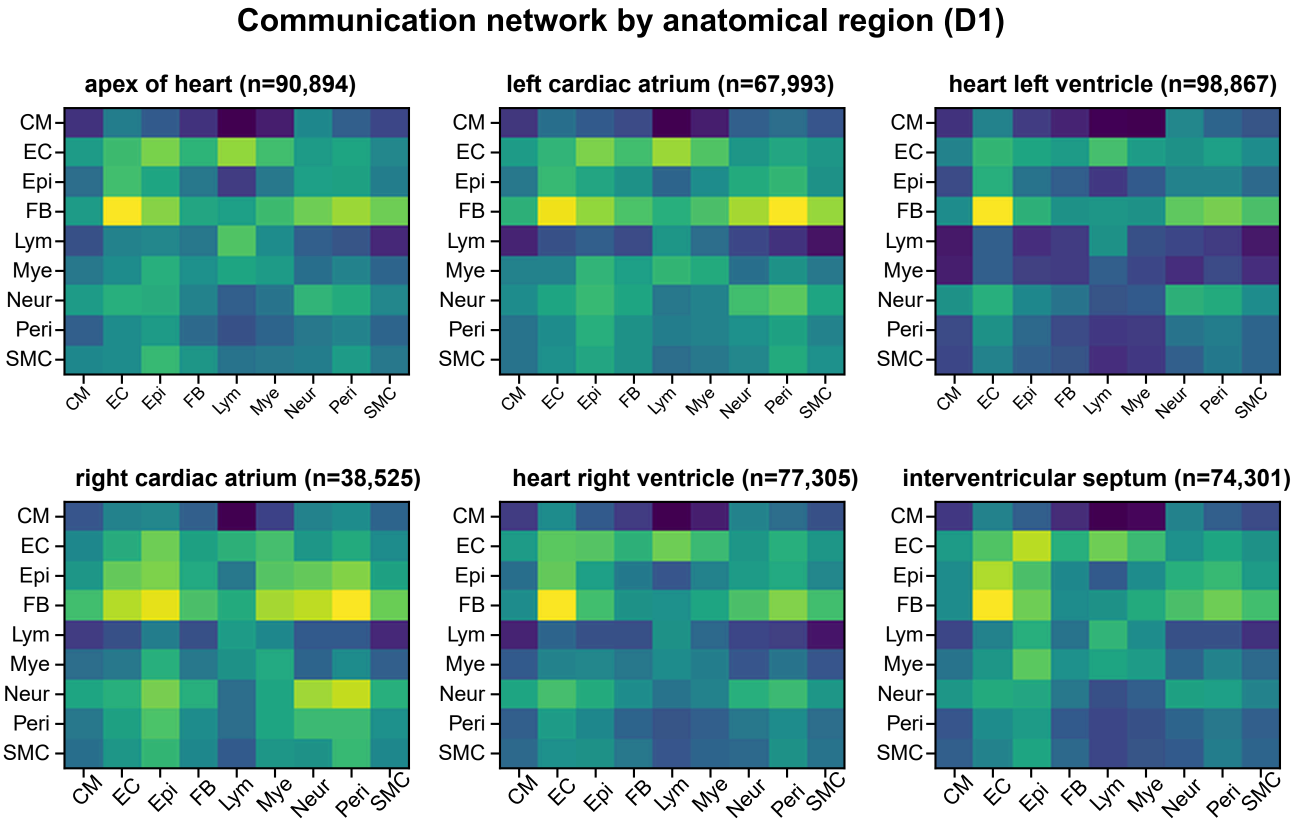


Figure S15. Region-specific communication networks. Ligand–receptor communication networks resolved for each anatomical region of the adult human heart.

#### Supplementary Figure S16

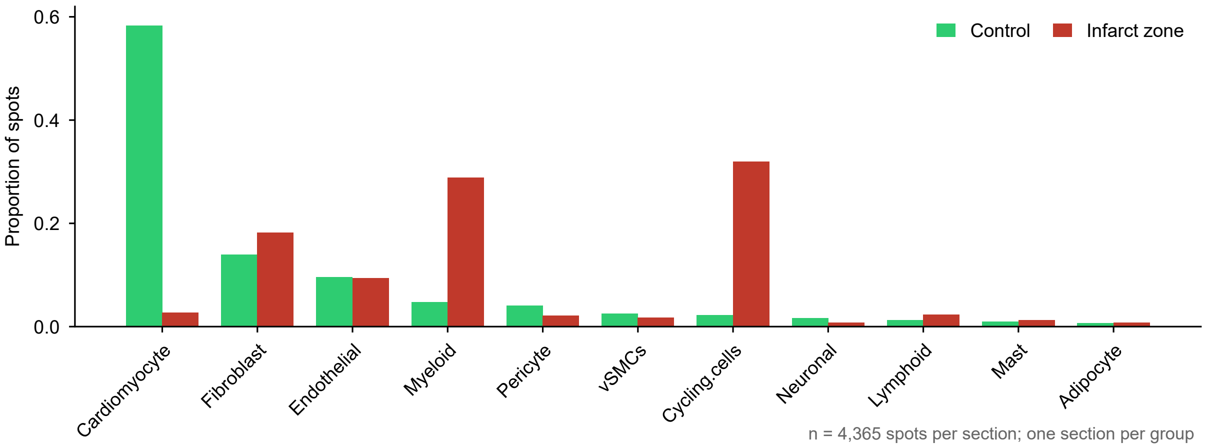


Figure S16. Deconvolved composition of infarct-zone and control Visium sections. Bars give the proportion of spots assigned to each cell class.

#### Supplementary Figure S17

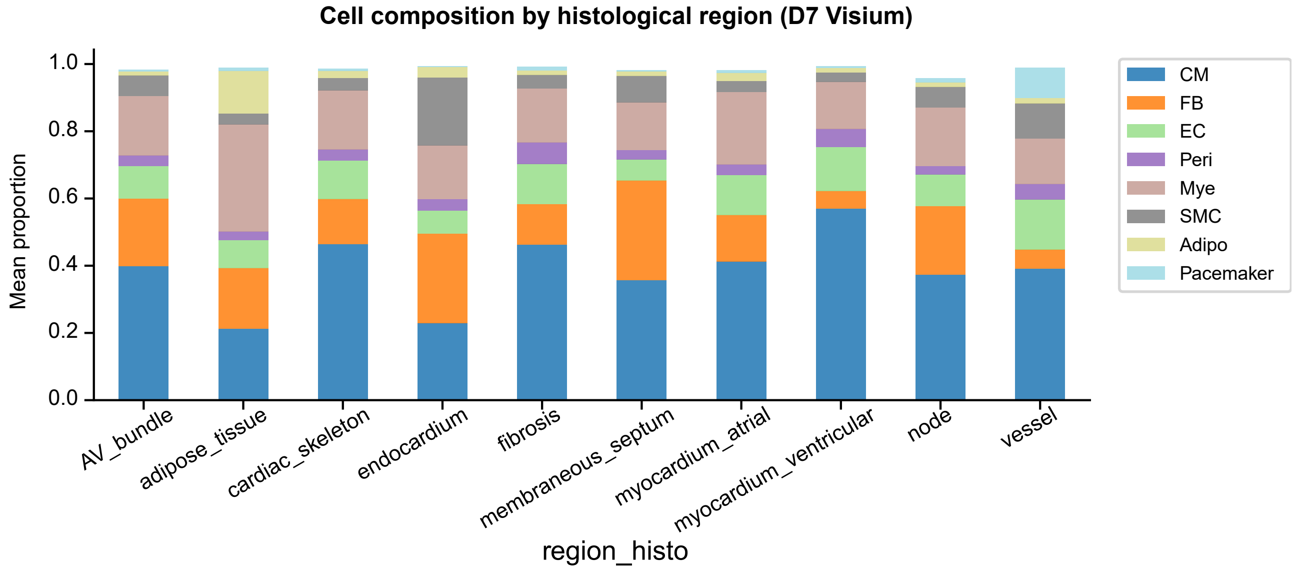


Figure S17. Histological-region cell-type composition. Deconvolved cell-type proportions across histological regions of the D7 Visium atlas.

#### Supplementary Figure S18

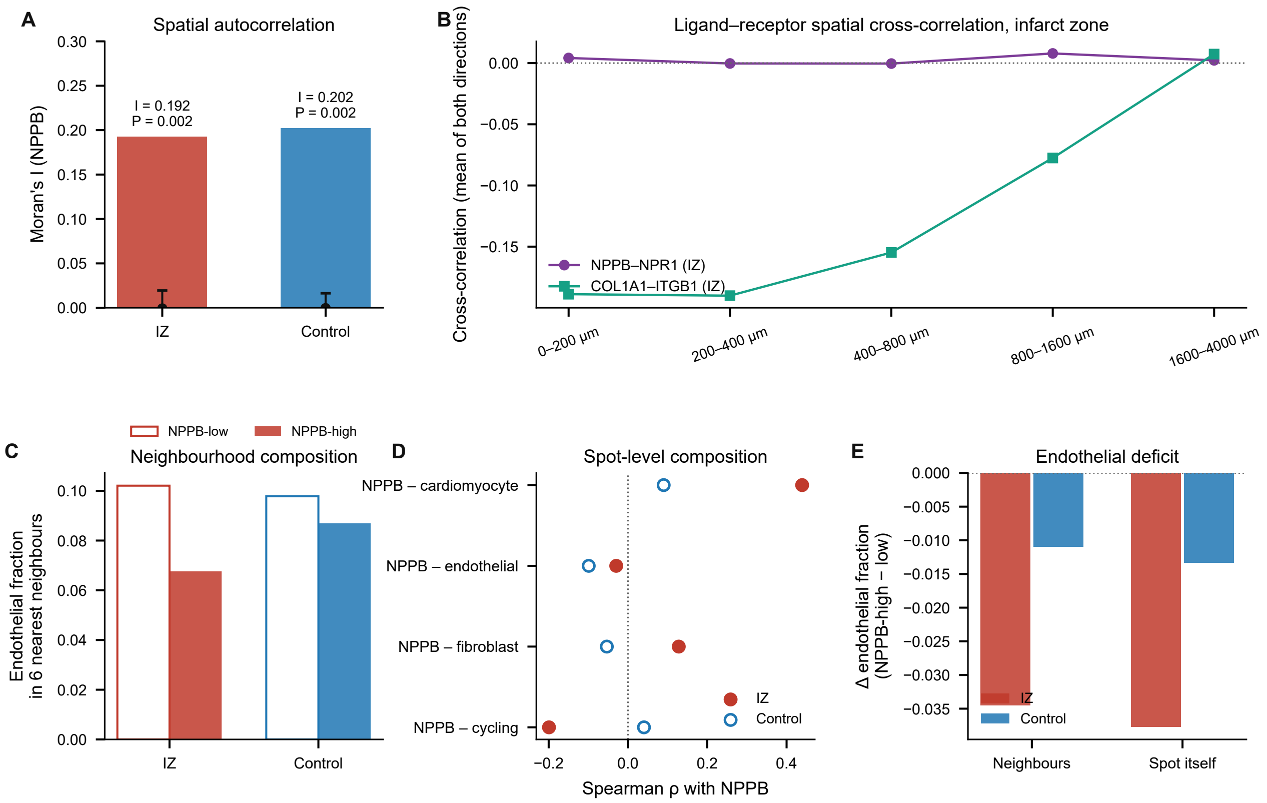


Figure S18. Spatial statistics of the natriuretic secretome. (A) Moran's I of NPPB with permutation nulls. (B) Distance-resolved cross-correlation between NPPB and NPR1 with COL1A1–ITGB1 as positive control. (C) Endothelial fraction in the six nearest neighbours of NPPB-high versus NPPB-low spots. (D) Spot-level correlations. (E) Endothelial deficit.

#### Supplementary Figure S19

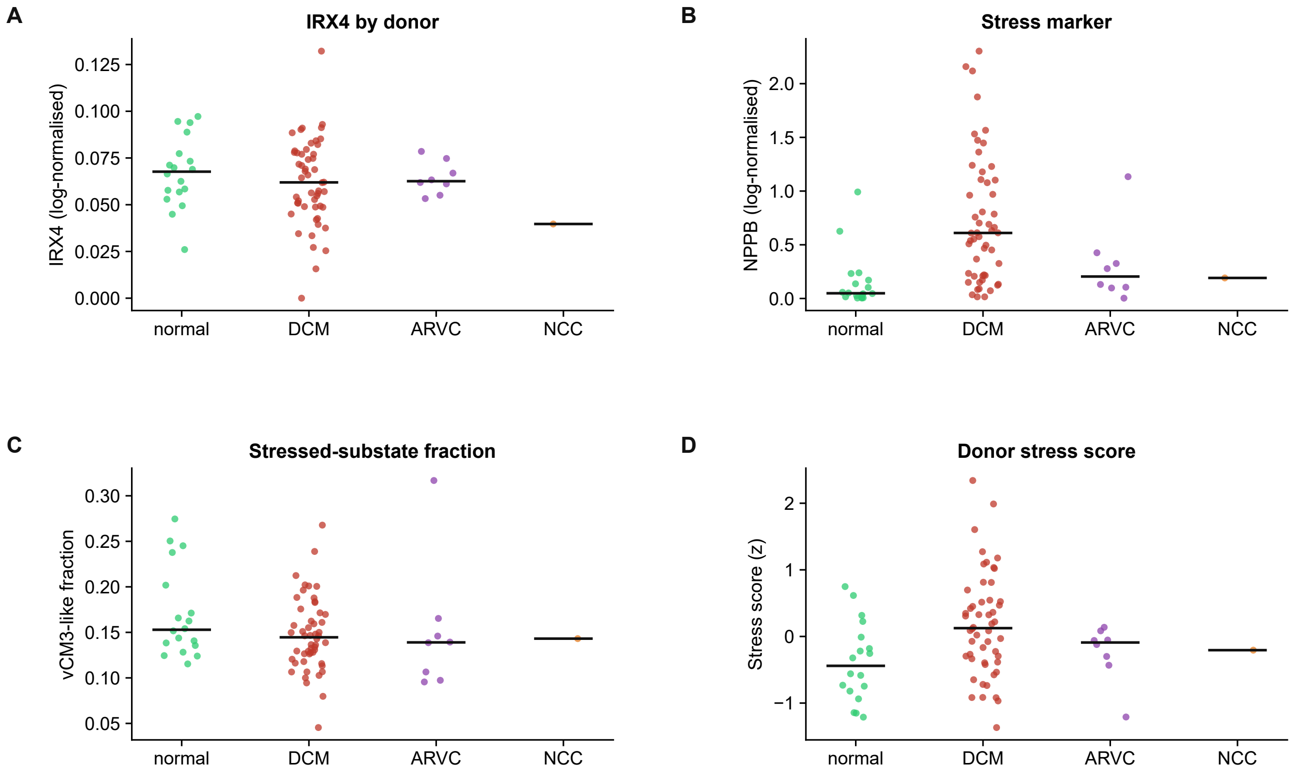


Figure S19. The independent cardiomyopathy cohort. (A) Donor-level IRX4 expression. (B) NPPB expression. (C) Fraction of vCM3_stressed-like cardiomyocytes. (D) Donor stress score. Groups are normal, dilated cardiomyopathy (DCM), arrhythmogenic right ventricular cardiomyopathy (ARVC) and NCC; each point is a donor and horizontal bars are medians.

#### Supplementary Figure S20

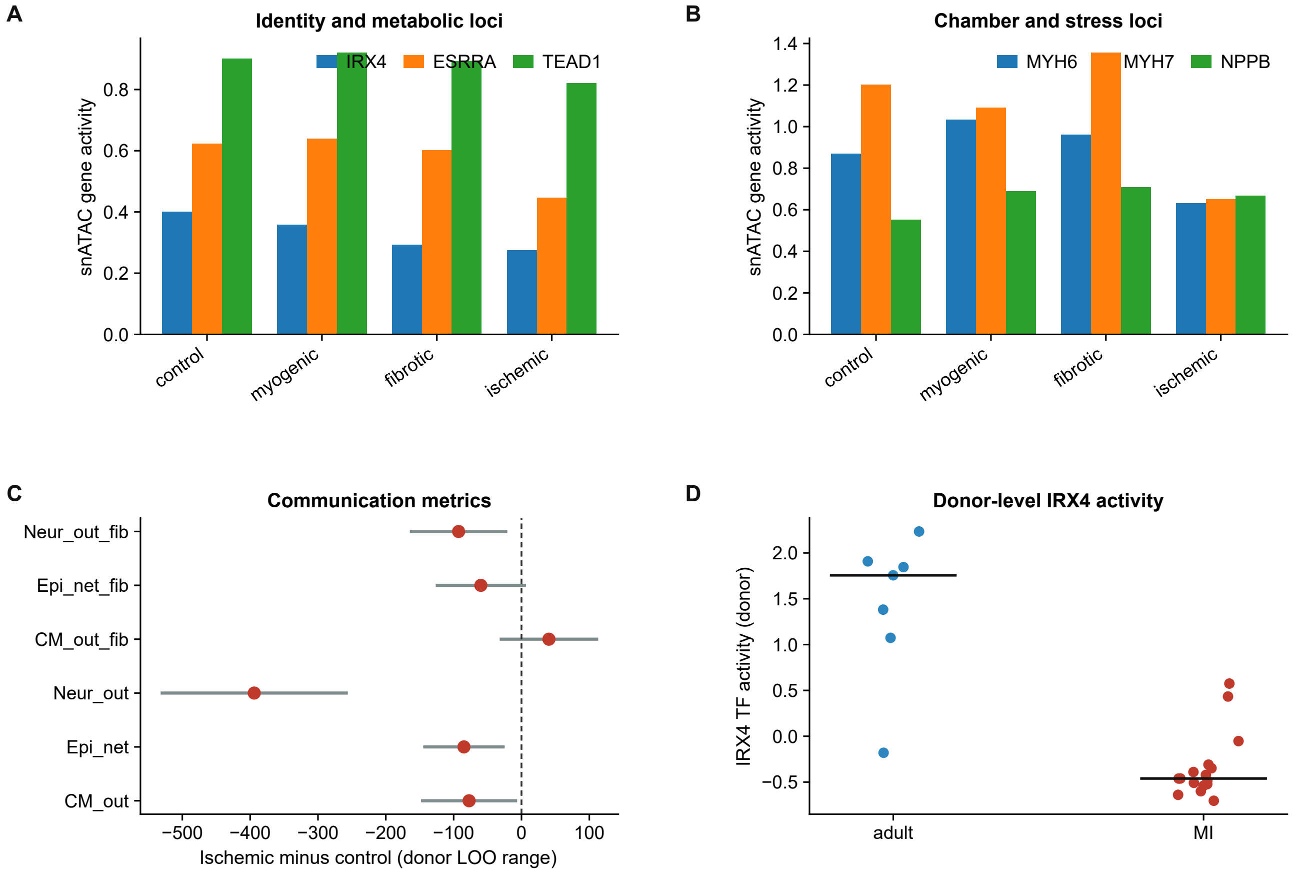


Figure S20. Zone stratification of the chromatin and communication layers. (A) snATAC gene-activity scores for IRX4, ESRRA and TEAD1 across control, myogenic, fibrotic and ischaemic zones. (B) MYH6, MYH7 and NPPB accessibility. (C) Communication metrics (ischaemic minus control, donor leave-one-out range). (D) Donor-level IRX4 activity in adult and infarcted myocardium.

### Supplementary Tables

Tables S1–S16 are provided as editable worksheets in the archive Supplementary_Tables_S1-S16.zip. Each table occupies one worksheet (Table_S1 to Table_S16), with supporting sub-sheets where indicated. The Index worksheet lists all sheets.

Table S1. Inventory of the public datasets and genetic resources used in this study.

Table S2. Programme scores versus developmental maturity.

Table S3. Marker expression, identity–stress scores and IRX4 pseudobulk across cardiomyocyte substates.

Table S4. Programme scores and donor-aware statistics in fetal, adult and infarcted cardiomyocytes; rare-event counts.

Table S5. Transcription-factor activity (decoupler, CollecTRI) across states and inference algorithms.

Table S6. Differential transcription-factor activity between fetal and adult cardiomyocytes.

Table S7. Single-nucleus ATAC gene-activity values, donor means and donor-clustered tests.

Table S8. Donor-level chromatin accessibility statistics and regulon enrichment.

Table S9. IRX4 locus scan in FinnGen release 13: lead variants, rs260399 effects and the quality-controlled heart-failure interval.

Table S10. cis-pQTL Mendelian randomization: instrument, harmonisation and outcome-specific estimates.

Table S11. Communication hub metrics for the D1 structural baseline.

Table S12. Cell-type by cell-type net communication strength in D1.

Table S13. D10 communication metrics, equivalence tests, null models and leave-one-out ranges.

Table S14. Deconvolved cell-type composition of the infarct-zone and control Visium sections.

Table S15. Spatial statistics for NPPB and the endothelial niche: Moran's I, distance-resolved cross-correlation and neighbourhood composition.

Table S16. Donor-level cardiomyopathy cohort statistics and aetiology-resolved expression.

#### Key Resources Table

| Reagent or resource | Source and identifier | Supplier |
|---|---|---|
| **Antibodies** | | |
| None used | This study did not use antibodies | — |
| **Bacterial and virus strains** | | |
| None used | This study did not use bacterial or viral strains | — |
| **Biological samples** | | |
| Public single-cell, single-nucleus and spatial transcriptomic datasets (D1–D10) | Dataset identifiers, cell numbers and original publications are given in Table S1 | CELLxGENE (cellxgene.cziscience.com) |
| **Chemicals, peptides, and recombinant proteins** | | |
| None used | This study used only public data | — |
| **Critical commercial assays** | | |
| None used | This study used only public data | — |
| **Deposited data** | | |
| FinnGen release 13 summary statistics (eight phenotypes) | https://r13.finngen.fi/ | FinnGen |
| deCODE plasma proteome pQTLs (NT-proBNP cis-pQTL rs198389) | ref. 26 | deCODE Genetics |
| Open Targets Platform release 26.09 | https://platform.opentargets.org/ | Open Targets |
| Analysis code | https://doi.org/10.5281/zenodo.22691010 (concept DOI) | Zenodo |
| **Experimental models: cell lines** | | |
| None used | This study did not use cell lines | — |
| **Oligonucleotides** | | |
| None used | This study did not use oligonucleotides | — |
| **Software and algorithms** | | |
| scanpy / anndata | 1.12.4 / 0.13 | https://scanpy.readthedocs.io |
| decoupler | 2.2.0 | https://decoupler.readthedocs.io |
| statsmodels | 0.15 | https://www.statsmodels.org |
| PyMuPDF / matplotlib | 1.26 / 3.11 | https://pymupdf.readthedocs.io |
| Custom analysis code | this paper | Analysis_Code_v31.zip (Supporting Information) |
